# *Plasmodium falciparum* impairs Ang-1 secretion by pericytes in a 3D brain microvessel model

**DOI:** 10.1101/2024.03.29.587334

**Authors:** Rory K. M. Long, François Korbmacher, Paolo Ronchi, Hannah Fleckenstein, Martin Schorb, Waleed Mirza, Mireia Mallorquí, Ruth Aguilar, Gemma Moncunill, Yannick Schwab, Maria Bernabeu

## Abstract

Disruption of the vascular protective angiopoietin-Tie axis is common in cerebral malaria (CM) patients, who display elevated angiopoietin-2 (Ang-2) and reduced angiopoietin-1 (Ang-1) blood concentrations. The role of pericytes in CM pathogenesis remains unexplored, despite being a major source of brain Ang-1 secretion and evidence of pericyte damage observed in CM postmortem samples. Here we engineered a human 3D microfluidics-based brain microvessel model containing the minimal cellular components to replicate the angiopoietin-Tie axis, human primary brain microvascular endothelial cells and pericytes. This model replicated pericyte vessel coverage and ultrastructural interactions present in the brain microvasculature. When exposed to *P. falciparum*-iRBC egress products, 3D brain microvessels presented decreased Ang-1 secretion, increased vascular permeability, and minor ultrastructural changes in pericyte morphology. Notably, *P. falciparum*-mediated barrier disruption was partially reversed after pre-treatment with recombinant Ang-1 and the Tie-2 activator, AKB-9778. Our approach suggests a novel mechanistic role of pericytes in CM pathogenesis and highlights the potential of therapeutics that target the angiopoietin-Tie axis to rapidly counteract vascular dysfunction caused by *P. falciparum*.

**The paper explained:** *Problem:* Cerebral malaria (CM) is a severe complication of *Plasmodium falciparum* infection, resulting in the majority of ∼600000 malarial deaths annually. Despite anti-malarial drug administration upon hospitalization, fatality rates still range from 15-25% and many survivors suffer long term neurological disabilities. A common dysregulated vascular pathway identified in CM patients is the angiopoietin-Tie axis. Treatments that restore this vascular homeostatic pathway appear as a potential avenue for adjunctive therapies in experimental rodent CM models. Nevertheless, the use of rodent CM models for therapeutic discovery is not ideal, given that *P. falciparum* pathogenesis is species-specific. Therefore, the development of novel and advanced human 3D microvascular models offers new avenues to study disease pathogenesis and explore potential adjunctive CM treatments.

*Results:* In this study, we generate a 3D human brain microvasculature model that reproduces *in vivo* interactions between two key cell types necessary to reproduce the protective angiopoietin-Tie axis: human brain endothelial cells and pericytes. Addition of *P. falciparum*-infected red blood cell (iRBC) egress products causes vascular disruption and hampers the release of the vascular protective factor, angiopoietin-1, from brain pericytes. 18-hour pre-treatment of Ang-1 for 18h prevents iRBC egress product-induced vascular disruption. A short pre-treatment of the microvessels with AKB-9778, a downstream pharmaceutical inducer of angiopoietin-Tie axis activity currently in phase II clinical trial for diabetic retinopathy, partially restores vascular integrity. Our study highlights the role of pericytes in CM and the therapeutic potential of interventions that restore the angiopoietin-Tie2 axis as adjunctive CM treatments.

*Impact:* Our study demonstrates the potential of bioengineered vascular models to recapitulate dysregulated pathways previously characterized in malaria patients, and in providing a physiologically-relevant platform to test adjunctive therapies. The use of the 3D brain microvascular model has enhanced our understanding of the mechanisms behind CM pathogenesis, uncovering a previously unappreciated effect of *P. falciparum* on brain pericytes, linking angiopoietin-Tie axis dysregulation and microvasculature disruption. These findings pave the way for the identification of novel, fast-acting therapeutics, such as AKB-9778, to restore vascular integrity in CM patients.

## Introduction

Cerebral malaria (CM) is a severe neurological complication of *Plasmodium falciparum* infection, clinically characterized by coma and fatality rates averaging 15-20% (Dondorp *et al*, 2010). Furthermore, approximately one third of CM survivors endure lifelong neurological sequelae such as hemiplegia, ataxia, epilepsy or speech disorders (Birbeck *et al*, 2010). The majority of pediatric CM-related deaths present severe brain swelling due to vasogenic edema, likely as a consequence of blood-brain barrier dysfunction (Mohanty *et al*, 2017). A hallmark of CM is the sequestration of *P. falciparum*-infected red blood cells (iRBC) in the brain microvasculature (Dorovini-Zis *et al*, 2011). While the molecular players responsible for *P. falciparum-*iRBC ligand-receptor interactions with cerebral endothelial cells have been extensively studied (Kessler *et al*, 2017; Storm *et al*, 2019; Sahu *et al*, 2022), the consequences of iRBC sequestration in the brain microvasculature require further investigation. Nevertheless, evidence suggests that increases in vascular permeability during CM are a result of a multifaceted process including the blockade of barrier-supportive endothelial receptors by iRBC, the release of toxic *P. falciparum* products from egressed iRBC, and a dysregulated host innate immune response (Bernabeu & Smith, 2017; Wassmer *et al*, 2024).

An important pathway commonly dysregulated in CM patients is the angiopoietin-Tie axis, crucial to maintaining endothelial pro-barrier, anti-inflammatory, and anti-apoptotic functions (Augustin *et al*, 2009). Binding of the ligand angiopoietin-1 (Ang-1) to the endothelial receptor Tie-2 promotes vascular quiescence and the stabilization of endothelial tight junction proteins, such as Occludin and Zona Occludens-1 (ZO-1), necessary for maintaining the barrier function of the brain microvasculature (Fukuhara *et al*, 2008; Saharinen *et al*, 2008; Siddiqui *et al*, 2015). Conversely, angiopoietin-2 (Ang-2), which is rapidly released from small endothelial storage granules called Weibel-Palade bodies, generally acts as a Tie-2 antagonist leading to endothelial activation and vascular leakage (Maisonpierre *et al*, 1997). Given the opposing roles of these two molecules in vascular function, increased levels of Ang-2 and decreased levels of Ang-1 have been largely associated with cerebral vascular pathogenesis in a multitude of diseases (Gurnik *et al*, 2016; Golledge *et al*, 2014). Similarly, a decrease in Ang-1, and an increase in Ang-2 and the Ang-2:Ang-1 ratio has been well documented in fatal cases of both pediatric and adult CM (Yeo *et al*, 2008; Lovegrove *et al*, 2009; Conroy *et al*, 2009, 2012; Jain *et al*, 2011). Despite its importance, the mechanisms leading to angiopoietin-Tie axis disruption in CM are poorly understood. Human pro- inflammatory cytokines and pro-coagulation proteins, such as TNF-α and thrombin, have been proposed to be responsible for the increase of Ang-2 secretion by endothelial cells (Gomes *et al*, 2023; Fiedler *et al*, 2004), but the causative agent responsible for decreased secretion of Ang-1 in CM patients remains unknown.

Vascular mural cells, including smooth muscle cells and pericytes are the main cellular sources for Ang-1 secretion in the brain microvasculature (Augustin *et al*, 2009). Beyond promoting endothelial barrier function through Ang-1 secretion, pericytes support the microvasculature and promote quiescence through additional roles. Structurally, pericytes wrap around cerebral microvessels and interact with the endothelium through reciprocal ultrastructural membrane protrusions, known as peg-and-socket junctions (Ornelas *et al*, 2021), and secrete extracellular matrix components of the basal lamina that strengthen microvascular architecture. Functionally, pericytes regulate brain blood flow by controlling vessel diameter through constriction (Daneman & Prat, 2015). Cerebral pericytes have been implicated in various vascular disorders of the central nervous system. Loss of pericytes from the cerebral vasculature during ischemic stroke and Alzheimer’s disease is associated with brain microvascular disruption, extravasation of blood components into the brain parenchyma, and neuronal loss (Fernández-Klett *et al*, 2013; Sengillo *et al*, 2013). Similarly, histopathological studies in retinal samples of fatal pediatric CM cases have revealed pericyte damage in regions of iRBC sequestration, suggesting a similar role of these cell types in severe malaria infections (Barrera *et al*, 2018). Yet, the role of cerebral pericytes in CM pathogenesis and in the dysregulation of the angiopoietin-Tie pathway remains largely unknown. Bioengineered vascular models are emerging as powerful tools to study disease mechanisms *in vitro.* They have been successfully used to recapitulate important vascular pathways, including the role of pericytes and the angiopoietin-Tie axis in the establishment of vascular networks (Haase *et al*, 2019), and more recently, malaria pathogenesis, by modeling ligand-receptor interaction of iRBC in microvessels (Bernabeu *et al*, 2019) or parasite-induced endothelial stress and inflammatory response (Howard *et al*, 2023). Here, we have developed a 3D brain microvascular model composed of primary brain endothelial cells and pericytes that recapitulates the minimal functional cellular unit of the angiopoietin-Tie axis found in the *in vivo* human brain. We have used this model to understand the interplay between iRBC, brain endothelial cells and pericytes that contributes to the molecular basis of angiopoietin-Tie dysregulation in CM. Our results indicate that products released during iRBC egress play a role in angiopoietin-Tie dysregulation by decreasing Ang-1 secretion by pericytes, highlighting the role of this cell type as a new player in CM pathogenesis. Furthermore, we showed that parasite-induced increase in endothelial permeability can be therapeutically targeted by the use of the angiopoietin-Tie axis inducers, such as the Tie-2 activator AKB-9778.

## Results

### *In vitro* 3D brain microvessel model recapitulates *in vivo* cerebral endothelial-pericyte interactions

Primary human brain microvascular endothelial cells (HBMEC) and human brain vascular pericytes (HBVP) were commercially purchased and the expression of cell type-specific markers was validated, including junctional proteins and transporters: vascular endothelial (VE)-cadherin, β-catenin, zonula occludens-1 (ZO-1), claudin-5 and glucose transporter-1 (GLUT-1), as well as, endothelial specific markers: CD31 and von Willebrand Factor (vWF) for HBMEC (Fig. EV1A,B). HBVP expressed brain pericyte markers platelet derived growth factor receptor β (PDGFRβ) and nerve/glial antigen 2 (NG-2) (Fig. EV1C) (He *et al*, 2016; Attwell *et al*, 2016). Furthermore, we confirmed HBMEC secretion of Ang-2 and Ang-1 exclusively secreted by HBVP in a transwell model (Fig. EV1D). To better recapitulate the brain microvascular architecture, we have generated a 3D brain microvessel model that recapitulates the minimal functional cellular unit of the angiopoietin-Tie axis. The device is fabricated in a collagen type I hydrogel containing a microfluidic network with a pre-defined geometry, which is connected to both an inlet and outlet, enabling vessel perfusion (Zheng *et al*, 2012). Previous incorporation of pericytes in similar bioengineered 3D models involved cellular addition into the collagen hydrogel itself. However, this resulted in sparse interaction of pericytes with the endothelial microvessels (Zheng *et al*, 2012). Nevertheless, *in vivo* cerebral pericytes have been shown to cover approximately one third of the brain microvasculature, albeit with slight variations along the brain vascular hierarchical network (Mathiisen *et al*, 2010; Uemura *et al*, 2020). To recreate this, HBVP were seeded directly into the channels along with HBMEC at a 5:1 endothelial to pericyte ratio (Fig. 1A). After 3 days in culture, the two cell types reorganized into two different layers with HBMEC forming a network of 100- μm diameter microvessels and mCherry-expressing HBVP covering the microvessel surface (Fig. 1B). HBVP distributed homogenously around the vessel cross-section, covering approximately a third of the microvessel surface area (29.6% of the bottom surface and 27.3% of the top surface) (Fig. 1C). Endothelial VE-cadherin junctional labeling is continuous along the vessel luminal surface, suggesting that the presence of HBVP does not disrupt the endothelial layer (Fig. 1D). Furthermore, HBMEC in the model retain the expression of endothelial adherens and tight junction markers, β-catenin and ZO-1 respectively (Fig. 1E). HBVP keep the expression of pericyte markers PDGFRβ and NG-2, as well as the mural cell contractile protein α-smooth muscle actin (αSMA), and appear as elongated cells with thin processes stretching over multiple endothelial cells (Fig. 1D, E). In addition, HBVP secrete laminin and collagen IV, contributing to the secretion of extracellular matrix components that compose the basal lamina of brain microvasculature *in vivo* (Fig. 1E) (Oliveira *et al*, 2023).

**Figure 1.**
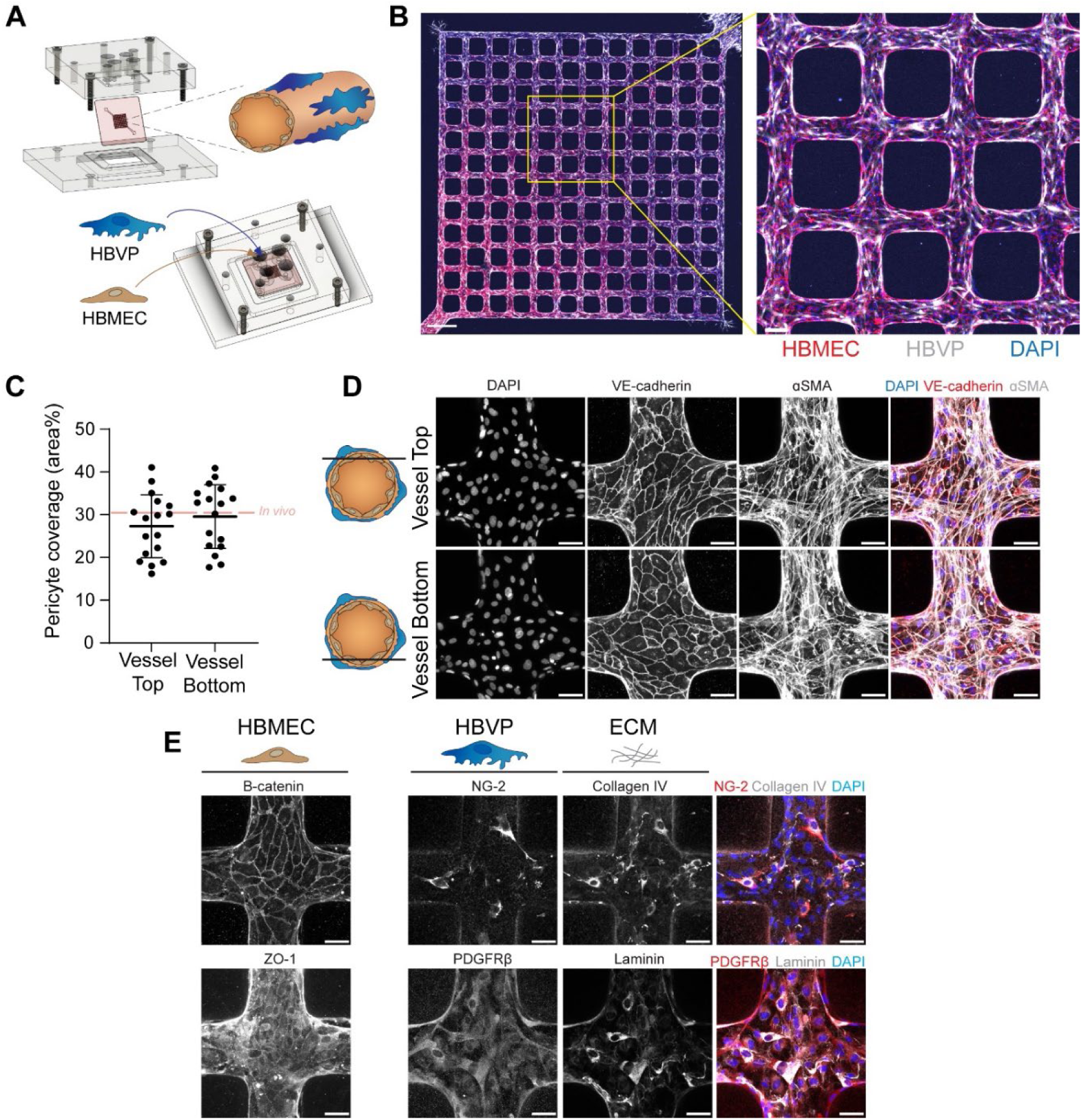
Generation and characterization of a 3D brain microvasculature model. **A** Schematic depiction of the fabrication pieces and device set-up (top-left), including a representation of resultant endothelial-pericyte interactions (top-right), and the seeding method used to generate the 3D microvessel model (bottom). **B** Immunofluorescence assay (IFA) maximum z-projection of the full network of cellularized channels labeled with Von-Willebrand factor for HBMEC (red), mCherry-expressing HBVP (white) and DAPI (blue) (left). Inset highlighting pericyte coverage of the microvessels (right). Scale bars: 500 μm and 100 μm (Inset). **C** Quantification of the percentage of endothelial microvessel area covered by pericytes on either the top or bottom microvessel surface in a z cross-sectional view. Red dashed line represents estimated *in vivo* brain microvascular pericyte coverage. Data is presented as mean +/- standard deviation (n = 17 devices, Mann-Whitney U test). **D** IFA maximum z-projection of VE-cadherin, αSMA, and DAPI labeling on the top and bottom cross-sectional surfaces of microvessels (left). Merged maximum z-projections with VE-cadherin (red), αSMA (white) and DAPI (blue) (right). Scale bar: 50 μm. **E** IFA maximum z-projection of brain endothelial markers, ZO-1 and β-catenin, as well as, colocalization between brain pericyte markers, NG-2 and PDGFRβ (red), and extracellular matrix markers, laminin and collagen IV (grey) in the presence of DAPI staining (blue). Scale bar: 50 μm.

Volumetric electron microscopy has been previously used to characterize *in vivo* 3D ultrastructural interactions between pericytes and endothelial cells of the cerebral microvasculature (Ornelas *et al*, 2021). To characterize the spatial and ultrastructural organization of pericytes and endothelial cells in the model, we utilized Serial Block-Face Scanning Electron Microscopy (SBF-SEM) (Denk & Horstmann, 2004), a volume electron microscopy technique that provides nanoscale resolution of samples in a large field of view (Peddie *et al*, 2022). We imaged a 100 μm segment of a microvessel branch with a 15x15x50 nm xyz resolution, followed by cellular segmentation and rendering. We characterized the spatial distribution and morphology of approximately 25 individual HBVP, distinguished from HBMEC by the presence of a highly granulated cytoplasm (Nahirney *et al*, 2016). Cross-sectional analysis and segmentation of the microvessel revealed an ovoid microvessel morphology, and confirmed the presence of HBVP at the abluminal side of the HBMEC (Fig. 2A). HBVP acquired a classical bump-on-a-log morphology, with the soma being the thickest region of the cell, and their longitudinal axis aligned with the direction of flow (Fig. 2B). Most of the HBMEC-HBVP interface is composed of thin pericytic lamellae (100-300 nm thickness) that cover large areas of the endothelial surface, with multiple thin branches appearing mostly at the pericyte periphery, as described previously (Ornelas *et al*, 2021). A close up of regions of HBVP-HBMEC cell interaction reveal that the membranes of both cell types are in close proximity (Fig. 2C). Pericytes in the brain constitute a heterogenous cell population depending on their location within the vascular branch. Considering the arteriole-sized diameter (100 μm) of the microvessels, the co- expression of αSMA, PDGFRβ and NG-2 staining, and the bump-on-a-log morphology, HBVP within the model are reminiscent of ensheathing pericytes, commonly found on cerebral arterioles rather than capillary pericytes that appear not to express αSMA (Smyth *et al*, 2018; Grant *et al*, 2019). Overall, the 3D *in vitro* brain microvessel model recapitulates *in vivo* interactions between brain endothelial cells and pericytes, both at the tissue and ultrastructural level, that can next be utilized to model the role of pericytes in malaria pathogenesis.

**Figure 2.**
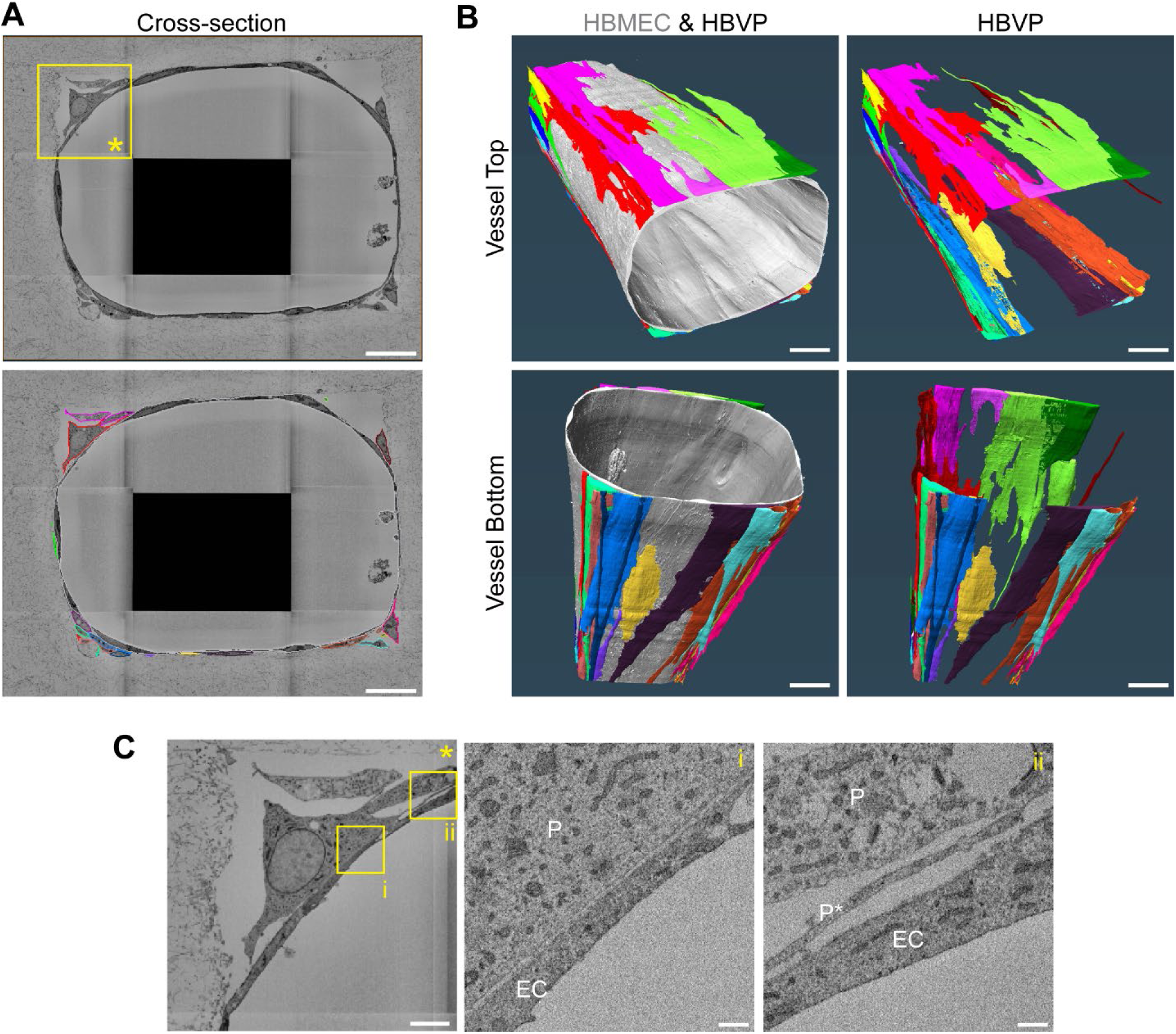
Serial block-face scanning electron microscopy reveals endothelial cell-pericyte ultrastructural interactions. **A** Cross-section of the microvessel imaged through the SBF-SEM volume (top) and the corresponding segmentation outlines of the endothelium (grey) and surrounding pericytes (coloured). Scale bar: 20 μm. **B** 3D rendering of the segmented microvessel (grey) and the surrounding pericytes viewed from above or below. Scale bar: 10 μm. **C** Zoomed images of the region of interest (ROI) in A. Insets show ROIs i and ii where pericytes are denoted with “P”, endothelial cells by “EC” and pericyte lamellae by “P*”. Scale bars: 5 μm (left) and 500 nm (center and right).

### *P. falciparum* egress products increase vessel permeability but only cause minor changes in pericyte morphology

*In vitro* studies have shown that *P. falciparum* products released during malaria parasite egress disrupt the endothelial barrier (Gillrie *et al*, 2007; Gallego-Delgado *et al*, 2016; Pal *et al*, 2016; Storm *et al*, 2019; Moxon *et al*, 2020). To characterize the impact that these could have on pericyte function, we established a protocol to create media enriched for *P. falciparum-*iRBC egressed products, denoted from here on as iRBC-egress media. Briefly, tightly synchronized schizont stage- iRBC were purified and allowed to egress in vascular growth media, followed by recovery of the supernatant fraction containing *P. falciparum* soluble products at an estimated concentration of 5x10^7^ ruptured iRBC/mL (equivalent to simultaneous egress of parasites at a parasitemia of approximately 1%) (Fig. EV2A, B). We measured the endothelial disruptive properties of iRBC-egress media by the xCELLigence system, which provides real time measurement of cell barrier integrity through impedance. Incubation with *P. falciparum* egress products caused a significant, dose-dependent decrease in cell index, a measurement of monolayer barrier function normalized to untreated cells, reaching a maximum disruption at 18 hours (Fig. EV2C). To elucidate if iRBC-egress media also induces barrier breakdown in the brain 3D microvessel model, we performed permeability assays in a simpler microvessel network which consists of a single 200 μm channel. After perfusion and incubation with iRBC-egress media for 18 hours, *P. falciparum* egress products accumulated on the walls of the microvessel channel and appeared as dark spots in brightfield suggesting the presence of parasitic hemozoin and food vacuoles (Fig. 3A). Microvessels perfused with iRBC-egress media presented a significant increase to 70 kDa FITC-dextran permeability across the entire vessel wall, corresponding to a 6-fold increment compared to vessels perfused with vascular cell media (4.08x10^-6^ cm/s in media-only vs. 2.66x10^-5^ cm/s in devices exposed to iRBC-egress media) (Fig. 3A, B). Although the presence of inter-endothelial gaps was not observed, VE-cadherin staining revealed areas of discontinuity and thinning of adherens junctions in regions nearby the deposition of parasitic DAPI-positive DNA remnants (Fig. 3C).

**Figure 3.**
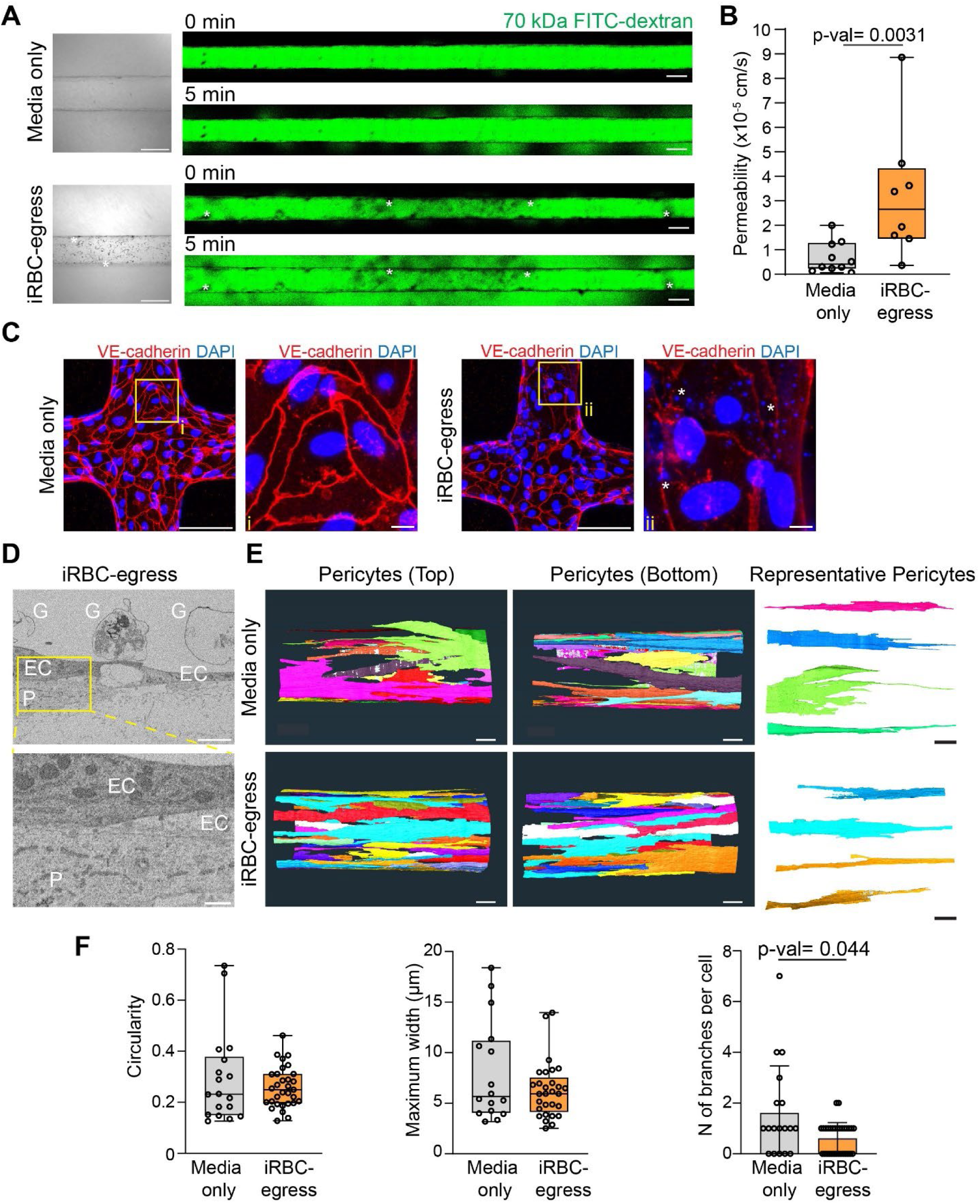
iRBC egress products increase microvessel permeability with minor changes in pericyte morphology. **A** Brightfield images of 3D brain microvessel channels after 18-hour incubation with media only or iRBC-egress media (left). Scale bar: 200 μm. Representative confocal images of 70 kDa FITC-dextran flux into the surrounding collagen 0 and 5 minutes after perfusion (right). Scale bar: 200 μm. Accumulated *P. falciparum-*iRBC egress material is represented with asterisks. **B** Apparent permeability of 70 kDa FITC-dextran in channels treated with media only or *P. falciparum-*iRBC egress media for 18-hours. (n = 8 individual devices). **C** Immunofluorescence maximum z-projection of 3D brain microvessels exposed to media only or iRBC-egress media stained with VE-cadherin (red) and DAPI (blue). Insets show ROIs of i and ii that display adherens junctions in the presence of cytoadhered DAPI-positive iRBC egress products represented by asterisks. Scale bar: 100 μm and 10 μm in insets. **D** SBF-SEM image and zoomed ROI of microvessels treated with iRBC-egress media. Pericytes are denoted as “P”, endothelial cells by “EC” and iRBC ghosts by “G”. Scale bar: 2 μm (top) and 500 nm in inset (bottom). **E** 3D rendering of pericytes segmented from microvessels treated with media only or iRBC-egress media and imaged with SBF-SEM (left). Scale bar: 10 μm. Representative segmented pericytes for morphology analysis (right). Scale bar: 10 μm. **F** Circularity, maximum width and number (N) of branches analysis (n = 18 or 31 segmented pericytes treated with media only or iRBC-egress media respectively). Data information: In **B** and **F** data are presented as box and whisker plots displaying the median, 25th and 75th percentiles and the minimum and maximum data points (Mann-Whitney U test).

To determine whether increased permeability was associated with physical changes in pericytes, we assessed their cellular morphology in 3D microvessels exposed to iRBC-egress media for 18- hours by SBF-SEM. HBVP maintained close spatial proximity to the endothelium in the presence of *P. falciparum*-iRBC ghosts, similar to control devices exposed to vascular growth media (Fig. 3D). To investigate potential changes in HBVP shape, we isolated and analyzed each 3D rendered HBVP (Figs. 3E,F and EV3A,B). Cellular features such as circularity and maximum width did not differ between conditions, however, iRBC-egress media treated HBVP presented fewer branches suggesting a response to iRBC-egress media (Fig. 3E,F). Altogether, our results show that *P. falciparum* egress products cause an increase in permeability of the 3D brain microvessel model, accompanied by minor changes in pericyte morphology.

### *P. falciparum-*iRBC induce dysregulation of the angiopoietin-Tie axis

Next, we aimed to determine whether parasite egress products are a major driver of angiopoietin- Tie axis disruption characteristic of CM (Yeo *et al*, 2008; Lovegrove *et al*, 2009; Conroy *et al*, 2009, 2012; Higgins *et al*, 2016). A multiplexed Luminex assay was used to measure secreted protein concentrations in microvessel supernatants recovered after 18-hour incubation with iRBC- egress media. Remarkably, we observed a substantial decline of Ang-1 secretion by HBVP following iRBC-egress media incubation (Fig. 4A). Consistent with recent studies on 2D HBMEC monolayers (Gomes *et al*, 2023), iRBC-egress media incubation did not result in increased Ang-2 secretion. Indeed, we quantified a significant decrease in Ang-2 secretion (Fig. 4A). Despite Ang- 2 decrease, a significant increase in the Ang-2:Ang-1 ratio was found (Fig. 4A). Additionally, we observed a decrease in 3D microvessel secretion or shedding of other vascular factors important for endothelial-pericyte interaction, such as Neural cadherin (N-cadherin) and Tissue Inhibitor of Metalloproteinase-1 (TIMP-1) (Fig.EV4A). Conversely, we detected no changes in soluble Vascular Endothelial Growth Factor (VEGF), Tie-2, Angiopoietin-like-4 (ANGPTL4), Platelet Derived Growth Factor-BB (PDGF-BB), Interleukin-8 (IL-8) or Chemokine Ligand-1 (CXCL-1) after 18-hour incubation with iRBC-egress media (Fig. EV4).

**Figure 4.**
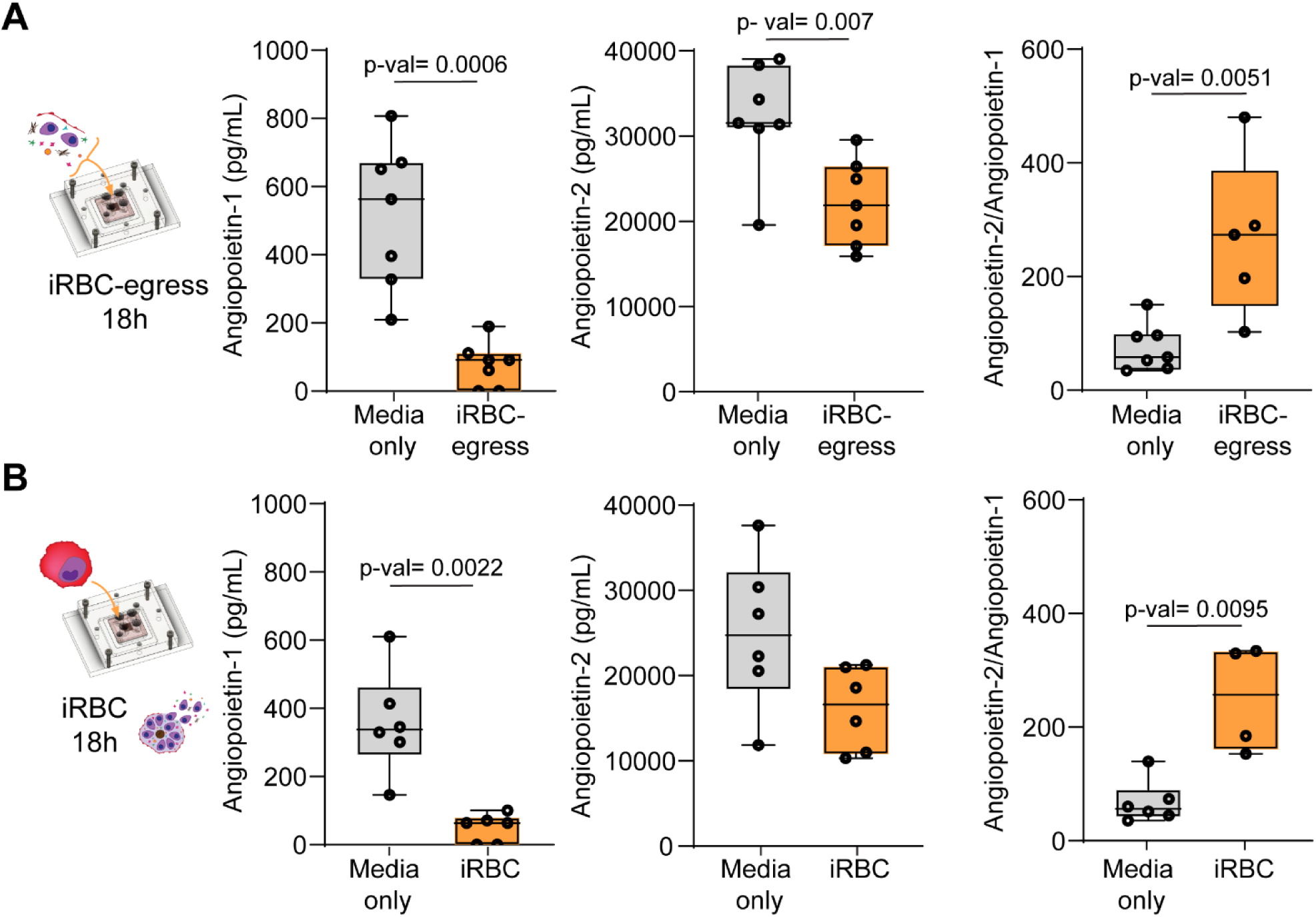
iRBC egress products induce angiopoietin-Tie axis dysregulation by altering pericyte secretion of Ang-1. **A, B** Ang-1, Ang-2 concentrations, and Ang-2: Ang-1 ratio measured from 3D brain microvessels treated with media only, iRBC-egress media (A) or *P. falciparum-*iRBC (B) for 18-hours (n = 6-7 pooled supernatants from 2-3 devices each) Data information: Box and whisker plots display the median, 25th and 75th percentiles and the minimum and maximum data points (Mann-Whitney U test) In **A** and **B**, data points in which Ang-1 concentrations were below the limit of detection were removed to calculate the Ang-2: Ang-1 ratio.

To validate whether changes in the angiopoietin-Tie axis occur physiologically following *P. falciparum* blood stage egress within the microvasculature, we perfused the 3D brain microvessels with schizont stage *P. falciparum-*iRBC for 30 minutes followed by a 10-minute wash. After an 18- hour incubation period, supernatants were again recovered and analyzed by Luminex. Similar results were obtained, with a reduction in both Ang-1 and Ang-2 secretion and an increase in the Ang-2:Ang-1 ratio (Fig. 4B). Altogether, these results suggest that *P. falciparum-*iRBC hamper Ang-1 secretion by pericytes, revealing a previously unappreciated role of pericytes in the dysregulation of the angiopoietin-Tie axis in CM.

### Therapeutic targeting of the angiopoietin-Tie axis partially protects against endothelial barrier breakdown by iRBC-egress media

Currently, there are no available host-targeted adjunctive therapies to restore cerebral vascular barrier function during CM. Given the key role of Ang-1 in promoting vascular quiescence and endothelial barrier formation we first investigated whether recombinant Ang-1 (rAng-1) supplementation could protect against the barrier disruptive effects of iRBC-egress media. First, we measured barrier disruption by xCELLigence in a 2D *in vitro* HBMEC culture in the absence of HBVP. An 18-hour pre-treatment of rAng-1 prior to HBMECexposure to iRBC-egress partially protected against parasite-mediated endothelial barrier disruption, while an 1-hour pre-treatment did not display a protective effect (Fig. EV5A-F). The 18-hour pre-treatment with rAng-1 conferred a 40% barrier breakdown protection after normalization to the untreated endothelial condition (Fig. EV5F). Next, we tested the protective capacity of rAng-1 against iRBC-egress products in the 3D microvessel model (Fig. 5A). Microvessels pretreated with rAng-1 did not present a significant increase in 70 kDa FITC-dextran permeability when exposed to iRBC-egress media. While vessels exposed to iRBC-egress media presented a median 6.4-fold increase in permeability, iRBC-egress vessels pre-treated with Ang-1 only presented a 1.3-fold increase in permeability compared to the rAng-1 treated control, equivalent to a 80% recovery in barrier breakdown (Fig. 5B,C). Taken together, pre-treatment with rAng-1 protects against the microvascular permeability caused by iRBC egress products, showing the potential for vascular restorative pathways that target the angiopoietin-Tie2 axis in CM treatment while highlighting the importance that a decrease in Ang- 1 could have in CM-induced vascular disruption.

**Figure 5.**
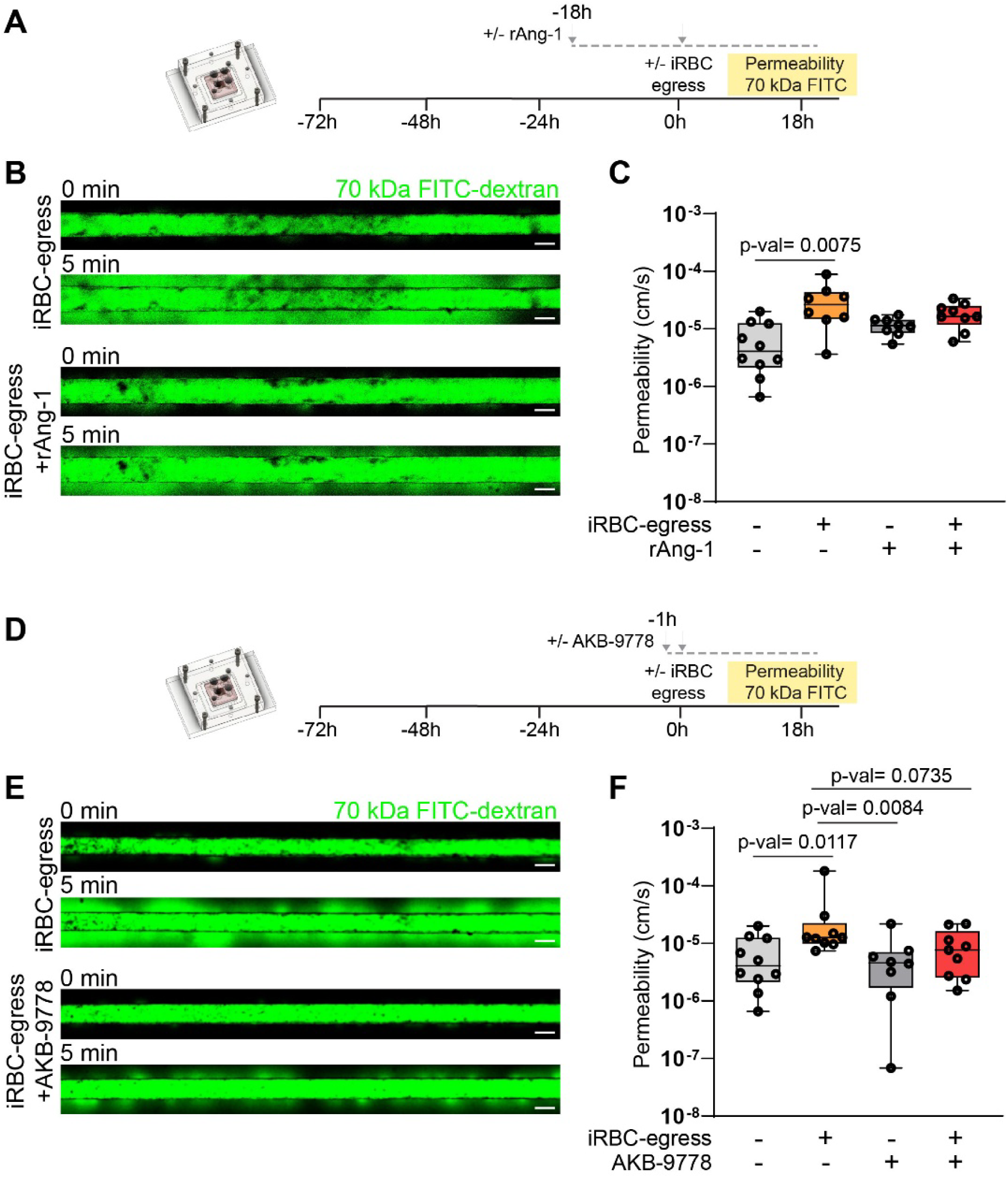
Therapeutic activation of the angiopoietin-Tie axis partially protects against increased permeability induced by iRBC egress products. **A** Experimental outline of the rAng-1 pre-treatment permeability experiment on 3D brain microvessels. **B** Confocal image of the 70 kDa FITC-dextran flux into the surrounding collagen at 0 and 5 minutes after dextran perfusion. Images correspond to a representative iRBC-egress treated microvessel pre-treated with either media only or rAng-1 for 18 hours. Scale bars: 200 μm. **C** Apparent permeability of 70 kDa FITC-dextran in microvessels pretreated with or without rAng-1 followed by 18-hour media only or iRBC-egress media treatment (n = 8-10 individual devices per condition. **D** Experimental outline of the AKB-9778 pre-treatment permeability experiment on 3D brain microvessels. **E** Confocal image of the 70 kDa FITC-dextran flux into the surrounding collagen at 0 and 5 minutes after dextran perfusion. Images correspond to a representative iRBC-egress treated microvessel pre-treated with either media only or AKB-9778 for 1 hour. Scale bars: 200 μm. **F** Apparent permeability of 70 kDa FITC-dextran in microvessels pretreated with or without AKB-9778 followed by 18-hour media only or iRBC egress media treatment (n = 8-10 individual devices per condition). Data information: Box and whisker plots display the median, 25th and 75th percentiles and the minimum and maximum data points in **C** and **F** (Kruskal-Wallis test corrected for multiple comparisons using the Benjamini, Krieger and Yekutieli method).

During angiogenesis, vascular endothelial-protein tyrosine phosphatase (VE-PTP) regulates Tie-2 activity by dephosphorylation (Fachinger et al., 1999). AKB-9778, a pharmacological inhibitor of VE-PTP activity, has been shown to specifically activate the angiopoietin-Tie axis to promote endothelial barrier function in the presence of various mediators of vascular disruption and inflammation (Shen et al., 2014; Frye et al., 2015). Therefore, AKB-9778 is currently undergoing phase II clinical trials as a therapeutic treatment for retinal vascular diseases characterized by angiopoietin-Tie axis dysregulation such as diabetic retinopathy (Sha et al., 2024). As the retinal and brain vasculature share similar embryological origin, anatomy, cell composition and response during CM (Patton et al., 2005; MacCormick et al., 2014), we tested the potential of AKB-9778 as a therapeutic option to counteract CM. In a 2D *in vitro* HBMEC culture, both 1- and 18-hour pre-treatment with AKB-9778 resulted in a partial protection against parasite-mediated endothelial barrier disruption of 56% and 43%, respectively (Fig. EV6A-F). Given the ability of AKB-9778 to protect against barrier disruption in 2D monolayers, we next validated its rapid protective activity in the 3D microvessel model (Fig. 5D). Microvessels pre-treated for 1h with AKB-9778 and exposed to iRBC-egress media presented a non-significant ∼2-fold increase in permeability while untreated microvessels presented a significant 3.2-fold increase in permeability, representing a 39% decrease in microvessel leakage after exposure to iRBC-egress media (Fig. 5E,F). The partial vascular barrier protective ability of AKB-9778 further highlights a crucial role of the angiopoietin- Tie axis in CM pathogenesis, as well as, suggests VE-PTP inhibition as a potential adjuvant therapeutic option for CM treatment.

## Discussion

Disruption of the angiopoietin-Tie axis is a common feature in CM patients, often associated with disease severity and fatality (Lovegrove *et al*, 2009; Conroy *et al*, 2012). This dysregulation is also found in pediatric and adult severe malaria patients without cerebral complications (Yeo *et al*, 2008; Conroy *et al*, 2009) or in placental malaria (Tran *et al*, 2021). Central to this pathway are brain pericytes, an Ang-1 secreting cell type that is located at the interface between the vasculature and brain parenchyma, yet largely understudied in the context of CM. Here, we developed a 3D brain microvessel microfluidic model that recapitulates the minimal functional brain microvascular unit to recreate the angiopoietin-Tie axis, including both primary brain microvascular endothelial cells and pericytes. Exposure of the 3D brain microvessels to *P. falciparum*-egress products caused an increase in vascular permeability accompanied by a decrease in Ang-1. Furthermore, we showed that pre-treatment with rAng-1 or the Tie-2 activator, ABK-9778, could partially protect against endothelial barrier breakdown. Taken together, these results suggest that the cessation of Ang-1 release upon exposure to iRBC-egress products is an important mechanism in microvessel disruption.

The bioengineered 3D model provides important vascular mechanical cues, including estimated flow and extracellular matrix stiffnesses present in the human brain. We could reproduce the high pericyte coverage found in the brain microvasculature by co-seeding primary brain endothelial cells and pericytes through the microfluidic network. Simultaneous cell seeding, at a ratio of 5:1 (endothelial cells:pericytes), allowed for the formation of an intact endothelial microvessel network with continuous labeling of adherens and tight junctional markers in the presence of pericytes with classical cerebral pericyte marker expression. SBF-SEM microscopy showed that both cell types sit at nanoscale distance, with the presence of thin lamellae extending from the pericyte soma that cover large areas of the endothelium. Overall, the 3D brain microvessel model recapitulates pericyte-endothelial interactions and coverage found in human brain vasculature making it suitable to study the role of pericytes in CM pathogenesis.

*P. falciparum* products released during blood stage egress have been shown to possess barrier breakdown capabilities on 2D endothelial monolayers grown on plastic. Proposed barrier disruptive *P. falciparum-*iRBC egress products include merozoite proteins, histones, heme, hemozoin and *P. falciparum* histidine rich protein 2 (Gomes *et al*, 2023; Gillrie *et al*, 2007; Gallego-Delgado *et al*, 2016; Pal *et al*, 2016; Storm *et al*, 2019; Moxon *et al*, 2020; Zuniga *et al*, 2022). However, their effects on pericytes remain unexplored. In this study, we confirmed that these products are disruptive in a microfluidics-based 3D brain microvessel model composed of primary human brain endothelial cells and pericytes. Pericytes play an important role in maintaining vascular homeostasis. For example, loss of pericytes from the cerebral vasculature leads to increased permeability in mouse models (Armulik *et al*, 2010). Furthermore, dysregulation of cerebral pericytes and detachment from the vasculature have been observed in other cerebral vasculature pathologies including stroke and Alzheimer’s disease (Fernández-Klett *et al*, 2013; Sengillo *et al*, 2013). In our study, we did not observe gross differences on pericyte morphology or loss of vascular coverage upon treatment with iRBC-egress media. Nevertheless, pericytes from iRBC-egress media*-*treated 3D microvessels did display subtle morphological changes, having significantly less branches. The relevance of these minor ultrastructural modifications in vascular barrier dysfunction could imply molecular mechanisms of disease. Luminex quantification upon stimulation with iRBC-egress media in the 3D brain microvessel model revealed a significant reduction in N- cadherin shedding. This could point towards reduced functional endothelial-pericyte interaction upon short-term incubation with *P. falciparum-*egress products. Future studies should explore whether exposure to *P. falciparum* for longer timepoints recapitulates phenotypes observed in post- mortem samples of cerebral malaria patients such as vacuolization (Pongponratn *et al*, 2003) or pericyte loss (Barrera *et al*, 2018). Nevertheless, our results point towards a loss of integrity at the endothelial and pericyte interface.

Ang-1 and Ang-2 have been considered in the past as potential biomarkers of CM (de Jong *et al*, 2016). Ang-1 has excellent predictive power to distinguish malaria severity scores (Conroy *et al*, 2009) or malaria from other central nervous system febrile diseases (Conroy *et al*, 2010). Nevertheless, the underlying mechanism of Ang-1 decrease in CM remains unknown, as previous *in vitro* studies overlooked the effect that iRBC could exert on brain pericytes. To our knowledge, our study is the first one to report that *P. falciparum* products released upon iRBC egress halt Ang- 1 secretion by pericytes. The presence of accumulated light scattering, DAPI-positive iRBC egress material suggests a role of extracellular parasitic DNA or hemozoin in the cessation of Ang-1 secretion, yet further studies are necessary to identify the specific iRBC egress component responsible for altered signaling pathways in pericytes. A complementary hypothesis for decreased Ang-1 serum concentrations in CM patients is severe thrombocytopenia, commonly associated with cerebral complications in *P. falciparum* infections (Sahu *et al*, 2022) since platelets are another major source of serum Ang-1 (Nachman & Rafii, 2008). In light of this, it is likely that both the effect of *P. falciparum-*iRBC on perivascular cells and thrombocytopenia could work in concert to further contribute to a widespread decrease of Ang-1 levels. Nevertheless, our results highlight brain pericytes as a new player in the development of CM through dysregulation in the secretion of Ang-1.

Clinical studies have shown a lack of correlation between Ang-2 levels and *P. falciparum-*iRBC microvascular obstruction in the rectal microvasculature (Hanson *et al*, 2015) or iRBC sequestration in brain post-mortem samples (Prapansilp *et al*, 2013). Similarly, our study confirms that *P. falciparum-*iRBC sequestration in 3D brain microvessel models does not directly cause Ang- 2 secretion by endothelial cells. This result is in concordance with a recent study that showed that Ang-2 is not secreted in response to *P. falciparum* lysate products, but rather occurs following endothelial activation with TNF (Gomes *et al*, 2023). Other studies have shown Ang-2 release as a result of exposure to thrombin or hypoxia (Pichiule *et al*, 2004; Fiedler *et al*, 2004), conditions that have also been associated with CM pathogenesis. Altogether, our study is in agreement with previous reports suggesting that *P. falciparum* sequestration does not play a direct role in the secretion of Ang-2, which instead is more likely a consequence of the activation of host pro-inflammatory factors.

Modulation of the angiopoietin-Tie axis as an therapeutic target against CM has long been of interest (Higgins *et al*, 2016) due to its role in strengthening endothelial tight junctions, as well as, upregulating anti-apoptotic and anti-inflammatory pathways (Saharinen *et al*, 2017). Interestingly, we showed that pre-incubation of human microvessels with recombinant Ang-1 protected against iRBC-egress mediated barrier breakdown. While the concentration of recombinant Ang-1 utilized in this study was supraphysiological compared to peripheral blood samples, it is quite likely that peripheral blood concentrations underestimate local concentrations at the endothelial-pericyte interface. Additionally, in our model, recombinant Ang-1 protection was only achieved after a pre- incubation of 18-hours, suggesting that downstream signaling through the Tie2 receptor, such as PI3K/Akt signaling, is required (Augustin *et al*, 2009).

Future interventions targeting the angiopoietin-Tie axis should focus on approaches that reverse microvascular dysfunction after exposure to *P. falciparum.* Pre-treatment with AKB-9778 resulted in rapid but partial protection against iRBC egress product-mediated breakdown, highlighting the potential of the angiopoietin-Tie axis as an adjuvant therapeutic target for CM treatment. Furthermore, AKB-9778-mediated activation of Tie-2 could provide an additional benefit in combatting CM pathogenesis as it has also been shown to block the binding of leukocytes to the vascular wall (Frye *et al*, 2015), with recent reports describing their accumulation in the brain vasculature of CM patients (Riggle *et al*, 2020). Previously, Tie-2 activators such as recombinant Ang-1 and the PPARγ agonist rosiglitazone have shown improved outcomes in a rodent experimental CM model after onset of disease, preserving blood-brain barrier integrity, and leading to a significantly increased survival rate (Higgins *et al*, 2016; Serghides *et al*, 2014). Phase II clinical trials elucidating the benefit of these therapeutics in human CM treatment are currently underway (Varo *et al*, 2024). Altogether, our results and others highlight the potential of compounds targeting the angiopoietin-Tie2 as adjunctive CM therapies.

To note, our study has several limitations, including the low-throughput of the model or challenges in growing the devices for long periods of time, hence preventing the modelling of long-term disruptive mechanisms, such as pericyte loss. Furthermore, our devices currently represent larger microvessels of the brain given the 100 μm diameter of our model and α-SMA expression on pericytes. Therefore, future studies could exploit 2-photon laser ablation or bioengineered self- assembled models (Arakawa *et al*, 2020; Hajal *et al*, 2022) to generate brain capillary-sized microvessels (5-10 μm), a major pathogenic site in CM. Nevertheless, this study establishes a key role of pericytes on microvessels that are similar in size to arterioles or post-capillary venules, regions of the brain vascular tree that are equally damaged in CM patients (Barrera *et al*, 2018). Finally, an additional limitation of our model is the absence of other cells that could play an important role in maintaining the angiopoietin-Tie axis. For example, Ang-1 secretion has been described in other brain and blood cell types, such as smooth muscle cells (Sundberg *et al*, 2002; Nishishita & Lin, 2004), platelets and more recently, neurons and astrocytes (Wei *et al*, 2021). Yet, one of the major advantages of *in vitro* bioengineered models is the opportunity to sequentially introduce different components that, independently or collectively, could play a role in a complex and multifaceted disease such as CM. Subsequent iterations of the model presented here could introduce other cell types that produce Ang-1 or incorporate pro-inflammatory cytokines or thrombin, to concurrently model functional consequences of increased secretion of Ang-2 by endothelial cells.

In summary, our study highlights the role that pericytes could have in the development of CM pathogenesis by showing for the first time that egress of *P. falciparum-*iRBC interrupts Ang-1 secretion by this cell type. In addition, our findings confirm that activation of the angiopoietin-Tie axis partially protects against increased vascular permeability mediated by *P. falciparum* egress products, providing further evidence that interventions involving this pathway could be part of a multi-targeted adjunctive therapy against CM.

## Materials and Methods

### Parasite lines

HB3var03 variant *P. falciparum* parasites (dual ICAM-1 and EPCR binding), regularly panned and monitored for appropriate PfEMP1 expression, were cultured in human B+ erythrocytes in RPMI 1640 medium (GIBCO) supplemented with 10% human type AB-positive plasma, 5mM glucose, 0.4 mM hypoxantine, 26.8 mM sodium bicarbonate and 1.5 g/L gentamicin. Parasites were grown in sealed top T75 flasks in a gas mixture of 90% N_2_, 5% CO_2_ and 1% O_2_. To maintain synchronous cultures, parasites were synchronized twice a week in their ring stage with 5% sorbitol and once a week in their trophozoite stage with 40 mg/mL gelaspan (Braun).

### P. falciparum-iRBC egress media preparation

Late-stage *P. falciparum-*iRBC (schizonts, parasitemia of 5-10%), synchronized in a 6-hour window, were purified by use of 40 mg/mL gelaspan gradient separation to a final purity of >60% parasitemia. The enriched *P. falciparum-* iRBC were then placed in complete RPMI media containing 1 μM compound 2, a reversible PKG inhibitor that inhibits iRBC egress (kindly donated by Michael Blackman, The Francis Crick Institute), at a concentration of 50 million *P. falciparum-*iRBC/mL. After 5 hours, compound 2 containing media was removed and the parasites were resuspended at a concentration of 100 million *P. falciparum-*iRBC/mL in vascular growth media, put into a sealed top T25 flask, gassed and left in the incubator overnight on a shaker set to 50 rpm to facilitate parasite egress. The resulting parasite egress efficiency was assessed by a hemocytometer count and blood smear, and was then concentrated to a working concentration of 50 million ruptured *P. falciparum-*iRBC/mL. The resulting iRBC-egress media is spun at 1000 rpm to remove cellular debris and flash frozen in liquid nitrogen until used. This concentration was chosen as this value is equivalent to a circulating parasitemia of 1%, concentrations frequently found in malaria patients.

### Primary human brain microvascular endothelial cell and primary human brain vascular pericyte culture

Primary HBMEC (Cell Systems; ACBRI 376) were cultured according to the manufacturer’s recommendations in complete endothelial growth media-2MV (Lonza) containing 5% fetal bovine serum. Cells were split using Trypsin/EDTA. When 90% confluent, HBMEC were seeded on 15 μg/mL Poly-L-lysine (P8920, Sigma) coated T75 flasks. Primary HBVP (ScienCell) were cultured according to the manufacturer’s recommendations in basal pericyte media supplemented with 1% pericyte growth supplement, 1% penicillin/streptomycin and 2% fetal bovine serum (ScienCell). Cells were split using Trypsin/EDTA when 90% confluent, seeded on 15 μg/mL Poly-L-lysine coated T75 flasks, and maintained in a humidified incubator at 37 °C and 5% CO_2_. All experiments were conducted with HBMEC and HBVP with a passage number of 7-9.

### mCherry lentivirus transduction

HBVPs were grown in a T75 flask to a ∼90% confluency and incubated with lentiviral particles containing a mCherry vector (kindly donated by Kristina Haase Lab, EMBL Barcelona) in serum free pericyte media at a multiplicity of infection of 10. After 24 hours, the lentivirus particles were removed by washing once every 24-hours for a 2-day period. mCherry-positive cells were then selected by fluorescence-activated cell sorting, expanded and froze down for future experiments.

### 3D brain microvessel model fabrication

Type 1 collagen was extracted from rat tails, dissolved in 0.1% acetic acid, lyophilized in a freezer dryer (Labconco Freezone 2.5 Plus) and resuspended at 15 mg/mL in 0.1% acetic acid for storage. The stock collagen solution-0.1% acetic acid solution was neutralized and diluted to 7.5 mg/mL on ice. A 13 by 13 channel grid or single channel pattern were negatively imprinted into the collagen hydrogel by soft lithography using a PDMS stamp, and inlet and outlets were created by insertion of stainless-steel dowel pins. The top pre-patterned collagen hydrogel contained in a plexiglass top jig is sealed to a flat collagen-layered bottom that sits on a coverslip and a plexiglass bottom jig. The assembly creates perfusable 120 μm diameter microvessels (grid design) or 200 μm diameter microvessels (single channel design), as described previously (Zheng *et al*, 2012; Piatti *et al*, 2022). Prior to cell seeding, the channels were incubated with vascular growth media, consisting in EGM-2MV (Lonza) supplemented with 1x pericyte and 1x astrocyte growth factors (ScienCell) for 1 hour. Primary HBMECs and HBVPs, or mCherry- positive HBVPs (for pericyte coverage analysis), were resuspended at concentrations of 7 million cells/mL and mixed to a ratio of 5:1 HBMECs to HBVPs. The cell mixture was then seeded into the inlet in 8 μL increments and driven through the microfluidic network by gravity-driven flow. Cells were perfused twice from either the inlet or outlet until the channels were completely covered with adhered cells. Media was then removed and devices were flipped upside down for 1-hour to ensure even cell distribution on the top and bottom of the channels. Microvessels were cultured for 3 days before being used in experiments with media change every twice per day by gravity driven flow.

### Immunofluorescence microscopy

Microvessel labeling was performed by gravity driven flow. Microvessels were fixed with 3.7% paraformaldehyde (PFA) in PBS for 15 minutes followed by three 10-minute washes with PBS. Next microvessels were incubated with Background Buster (Innovex) for 30 minutes and then permeabilized with blocking buffer (0.1% Triton X-100 and 2% bovine serum albumin in phosphate-buffered saline (PBS)). Primary antibodies including rabbit anti-αSMA-555 (EPR5368, Abcam), mouse anti-vWF (sc-365712, Santa Cruz), mouse anti-ZO-1 (Invitrogen, 10017242), rabbit anti-β-catenin (Cell Signaling, 9587S), mouse anti-PDGFRβ (Abcam, ab69506), mouse anti-NG-2 (Invitrogen, 10424493), rabbit anti-laminin (Abcam, ab11575), rabbit anti-collagen IV (Abcam, ab6586)), mouse anti-VE-cadherin (sc-52751, Santa Cruz), and Phalloidin-647 (Invitrogen, A22287) were diluted in blocking buffer at 1:100 and incubated overnight at 4 °C. After three 10-minute PBS washes, secondary antibodies including goat anti-mouse Alexa Fluor 488 (A11001, Invitrogen), goat anti-rabbit Alexa Fluor 488 (Invitrogen, A-11008), goat anti-mouse Alexa Fluor 594 (Invitrogen, A-11005), goat anti-rabbit Alexa Fluor 594 (Invitrogen, A-11012), goat anti-mouse Alexa Fluor 647 (A21235, Invitrogen), goat anti-rabbit Alexa Fluor 647 (Invitrogen, A-21244) and 2 mg/mL DAPI (D21490, Invitrogen) were diluted at 1:250 in 2% bovine serum albumin and 5% goat serum containing PBS, and incubated at room temperature for 1 hour. Microvessels were washed 6 times with PBS for 10 minutes each and then imaged on a Zeiss LSM 980 Airyscan 2. Image stacks were acquired with 3 μm or 10 μm z-step size for single or tile scan images respectively. Z-projections and further threshold analysis was done using Fiji (ImageJ v1.54f) software.

#### Pericyte coverage analysis

Tile scan images of the microvessel network with mCherry-positive HBVP and vWF-labeled HBMEC were acquired as described above. Images were divided into two stacks, one being the top half of the microvessels of the network and the other the bottom half. Next, Z-projections of the top and bottom surfaces of the network were generated. The percentage of endothelial surface that was covered by pericytes was calculated by creating a mask of both the mCherry HBVP and the vWF-positive HBMEC.

#### Serial Block-Face Scanning Electron Microscopy

Microvessel devices were grown for 3 days, then treated with iRBC-egress media or vascular growth media for 18hours, and then fixed by adding 2% PFA and 2.5% glutaraldehyde (GA) to the device inlet for 30-minutes and washed twice with EGM-2MV at 37 °C. The collagen hydrogel was then carefully removed from the plexiglass jig and microvessel regions exposed to low shear stress were cut out and fixed with a secondary fixative solution (2% PFA, 2.5% GA, 0.25 mM CaCl_2_, 0.5 mM MgCl, 5% sucrose in a pH 7.4 0.1 M Cacodylate buffer) overnight at 4 °C, and then rinsed twice for 15-minutes with 0.1 M Cacodylate buffer. Samples were post-fixed in a reduced Osmium solution (1% OsO_4_, 1.5% K_3_FeCN_6_ in 0.065 M Cacodylate buffer) for 2-hours at 4 °C followed by six 10-minute washes in dH_2_O. Post-staining consisted of subsequent incubation steps of 1% thiocarbohydrazide in dH_2_O, 2% Osmium tetroxide in dH_2_O, and 1% Uranyl acetate in dH_2_O, aided by a PELCO Biowave Pro+ (Ted Pella) containing a SteadyTemp Pro and ColdSpot set to 20 °C at 7x 2 minute cycling on-off at 100 W under vacuum, with dH_2_O rinses in between steps once in a fume hood and twice in the microwave at 250 W for 40 s without vacuum. The stained samples were then dehydrated by serial additions of 30, 50, 80, and 100% ethanol solutions in the microwave at 250 W for 40 s without vacuum with ColdSpot set to 4 °C, infiltrated with EPON 812 hard epoxy resin in steps of 25, 50, 75, 90, and 3x 100% resin diluted in ethanol with each step in the microwave at 150 W for 3 minutes under vacuum and polymerized at 60 °C for 48 hours. The sample was trimmed (UC7, Leica Microsystems) using a 90° cryo-trimmer (Diatome) to generate a small block face. The resulting resin block was mounted on a pin stub using silver conductive epoxy resin (Ted Pella). The SBF-SEM acquisition was performed with a Zeiss Gemini2 equipped with a Gatan 3view microtome and a focal charge compensation device (Zeiss). The SEM was operated at 1.5 kV 300 pA, using a pixel size of 15 nm, a slice thickness of 50 nm and a dwell time of 1.6 μs. The SBF-SEM acquisition was performed using the software SBEMimage (Titze *et al*, 2018). The acellular center of the lumen was not imaged to save imaging time. Following acquisition, the positional metadata of the sections and tiles obtained was converted with SBEM-specific import scripts (https://git.embl.de/schorb/rendermodules-addons) for subsequent assembly, stitching and alignment of large EM datasets (Mahalingam *et al*, 2022). Aligned images were then binned to 22 by 22 by 250 nm^3^ in xyz and segmentation was then performed using the software package Amira (Thermo Fischer Scientific). Primary HBVP and HBMEC were segmented semi manually using the magic wand and brush tool to segment every fifth section followed by use of the interpolation tool to segment the entire volume. The resulting segmented microvessels were rendered using the generate surfaces module for 3D visualization.

#### P. falciparum-iRBC and iRBC-egress media perfusion to 3D brain microvessels

All experiments were done in 3D microvessel devices grown for 3 days. Schizont-stage *P. falciparum-*iRBC were purified to >60% parasitemia by 40 mg/mL gelaspan and the resulting enriched population was diluted in vascular growth media to 50 million iRBC/mL (same concentration as the *P. falciparum-* iRBC-egress media). 200 μL of *P. falciparum-*iRBC or *P. falciparum-*iRBC-egress media was added into the device inlet and perfused by gravity flow for 30-minutes with the outlet effluent being reintroduced to the inlet every 10 minutes. Microvessels treated with intact *P. falciparum-* iRBC (but not *P. falciparum-*iRBC-egress media), were then washed for 10 minutes with vascular growth media. Both conditions were incubated overnight for 18 hours. Supernatant was then removed from the outlet and pooled for Luminex analysis, and 3D brain microvessels were used for permeability studies or fixed for imaging with 3.7% PFA for 15-minutes (see section immunofluorescence microscopy).

#### Fluorescent dextran-based permeability assay

Single channel microvessel devices were grown for 3 days, then perfused and incubated for 18 hours with either vascular growth media or *P. falciparum-*iRBC-egress media as described above. To determine resulting changes in permeability, they were perfused with a 70 kDa FITC-dextran solution as follows: the device was washed 1x with PBS and placed in a Zeiss LSM 980 Airyscan2 confocal microscope with a temperature and CO_2_ controlled imaging chamber (37 °C, 5% CO_2_). PBS was aspirated and 100 μL of 70 kDa FITC-dextran solution (100 μg/mL) in PBS was pulled from the inlet to the outlet at 6.5 μL/min flow rate using a syringe pump (Harvard Apparatus PHD 2000). Tilescan confocal images of the entire channel were taken every 30 seconds for 5 minutes once the channel had been filled with FITC dextran. In devices pre-treated with 1 µg/mL recombinant Ang-1 (923-AB-025, R&D Systems), pre-incubation occurred at 18-hours, starting on day 2. In devices pre-treated with 10 µM AKB-9778 (HY-1009041-1MG, MedChemExpress), pre-incubation occurred 1 hour prior to iRBC-egress media or media addition.

#### Quantification of apparent permeability

Apparent permeability is determined as the flux of fluorescently labeled dextran across the microvessel wall into the surrounding collagen hydrogel. All steps of permeability analysis were performed using ImageJ. By using the following equation:

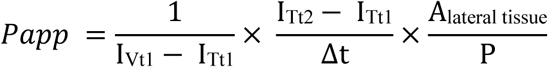

The apparent permeability (P_app_) was calculated using the fluorescence intensity inside the vessel (I_V_) and in the surrounding collagen hydrogel (I_T_) at two time points t1 (0 minutes after the vessel is filled with dextran) and t2 (5 minutes after the vessel is filled). Δt is the change in time, A_lateral tissue_ is the area of the collagen hydrogel being examined and P is the perimeter of the microvessel. To account for *P. falciparum-*iRBC-egress material that appears dark even in the presence of 70 kDa FITC-dextran, regions of *P. falciparum-*iRBC-egress material were excluded from the calculation of the fluorescence intensity inside the vessel (I_V_) by using ImageJ median smoothing and the magic wand tool to select only the area of the channel without *P. falciparum*-iRBC-egress material.

#### Serial Block-Face Scanning Electron Microscopy pericyte morphology analysis

3D-rendered pericyte meshes, segmented from SBF-SEM images of microvessels treated with either media alone or iRBC-egress media, were generated using Amira. Following segmentation, the meshes were exported as *STL* files and processed with a custom-developed Python script, designed to quantify aspects of pericyte morphology, including mesh circularity and maximum width (waleedmirzaPhD, 2024). Firstly, the 3D pericyte meshes were mapped onto a 2D plane through principal component analysis (PCA), effectively reducing the data’s dimensionality while retaining its most significant morphological features. This mapping uses the two principal components that encapsulate the dimensions with the maximum variance in the 3D mesh, thus ensuring the conservation of structural information. Subsequently, convex hull analysis (Virtanen *et al*, 2020) was applied to delineate the pericyte mesh boundaries, providing a clear representation of each structure’s shape by which circularity and maximum width could be measured. The circularity of each mesh is calculated using the convex hull, by assessing the ratio of the area enclosed by the hull to the square of the hull perimeter, with the formula for circularity given as 4𝜋 × *Area*/*Perimeter*^2^ . This circularity metric provides insight into the roundness of the mesh, where a value closer to 1 indicates a shape that is more circular, and values closer to 0 reflect more elongated or irregular shapes. Additionally, the maximum width is defined by identifying the two farthest points along the convex hull boundary and then calculating the greatest perpendicular distance from this line (connecting the farthest points) to any point within the convex hull. This measurement represents the maximum width of the pericyte mesh by assuming its length is the axis with the greater size. In addition, the number of pericyte branches was calculated manually by counting the number of branches on each 3D rendered pericyte.

#### Co-culture transwell fabrication

10^5^ HBVPs were seeded on the basolateral side of Poly-L-lysine treated transwell inserts (12 well PET membrane inserts with a pore size of 3 μm) in 100 μL of media and left to attach for 4-6 hours. Transwell inserts were then flipped into a 12 well culture plate containing 1200 μL of pericyte media and allowed to grow for 2 days. Next, 10^5^ HBMECs were seeded onto the apical side of the Poly-L-lysine treated transwell inserts in 300 μL of media and left to attach for 4-6 hours. Then media was removed from both sides of the insert and 1200 μL of pericyte media was added to the basolateral side and 800 μL of EGM-2MV media to the apical side. The cells were allowed to grow for 2 more days. On day 5, supernatants were taken from the basolateral side for angiopoietin-1 ELISA. Endothelial cell-only transwells underwent the same fabrication protocol minus the initial pericyte steps.

#### Angiopoietin-1 ELISA

Ang-1 protein concentrations were measured from supernatants taken from the basolateral side of the transwell model. Supernatant concentrations were obtained by running 5X diluted samples on either the Human Angiopoietin-1 DuoSet ELISA kit (DY923, R&D Systems). Protein concentrations were interpolated from sigmoidal 4-parameter-fit standard curves generated from a 7-point standard curve of recombinant human proteins using GraphPad Prism.

#### Quantification of secreted proteins by Luminex Assay

Secreted protein concentrations were measured from supernatants taken from the outlets of 3D brain microvessels with a 13 x 13 grid geometry. Supernatants from 2-3 devices exposed to the same condition were pooled, diluted 2X and assayed using a 10-plex Human Luminex Discovery Assay from R&D Systems on a Luminex 100/200. The 10-plex panel included: Ang-1, Ang-2, PDGF-BB, N-cadherin, TIMP-1, ANGPTL4, VEGF, Tie-2, IL-8, and CXCL-1. Protein concentrations were interpolated from a 11-point standard curve of known concentrations of recombinant human proteins provided by the vendor and reported as pg/mL using xPONENT 4.2. 7 out of 10 analytes were in the standard curve quantification range for 100% of measurements recorded, with Timp-1, N-cadherin and IL-8 having 100% of measurements above the upper limit of quantification.

#### Measuring changes in barrier integrity by xCELLigence

96 well PET E-plates (300600910, Agilent) were coated with 15 μg/mL Poly-L-lysine and 5000 HBMECs in 200 μL EGM-2MV were added per well. After cell adherence, the plate was placed into the xCELLigence RTCA SP reader to begin baseline measurement of growth-related changes in cell index (an arbitrary measure of impedance) and the cells were incubated for 3 days until cell index reached a plateau (indicative of cell confluence). Media was changed every two days. For initial testing of the impact of iRBC- egress media on barrier integrity, on the day of the experiment iRBC-egress media was added at concentrations of either 12.5, 25 or 50 million ruptured iRBC/mL. Changes in cell index were measured every 15-seconds for 8 hours and then every 15-minutes for 48-hours. To test the impact of rAng-1 or AKB-9778 on barrier breakdown mediated by *P. falciparum-*iRBC-egress, iRBC- egress media was added at 50x10^6^ ruptured iRBC/mL after either a 1- or 18-hour pre-incubation with 1 μg/mL of rAng-1 or 10 µM of AKB-9778. Again, changes in cell index were measured every 15 seconds for 8 hours and then every 15-minutes for 48-hours. Quantification of partial protection was done by first normalizing each condition to the media only control. Then the area under the curve of the negative values, indicative of barrier breakdown, was calculated using GraphPad Prism (version 10.0.2) where the media only without rAng-1 condition represented an average value of 0 and the iRBC-egress media without rAng-1 condition an average value of 100.

#### Statistical analysis

All statistics were obtained by use of GraphPad Prism (version 10.0.2). Non-parametric Mann-Whitney U tests were performed for most experiments, except to calculate Ang- 1 protection in the 2D xCELLigence and microvessel experiments in which a repeated measures one-way ANOVA test with Dunnett’s multiple comparisons test and Kruskal-Wallis test with Dunn’s multiple comparisons test were performed respectively. A P-value<0.05 was considered statistically significant.

## Author contributions

R.L and M.B conceived the work. R.L, P.R, R.A, and M.B designed experiments. R.L performed most experiments with assistance of F.K, P.R, H.F, M.M, and R.A. R.L, P.R, H.F, W.M, R.A, Y.S, G.M, M.B analyzed the data. R.L and M.B wrote the original draft of the manuscript. All authors contributed to manuscript writing, revision, editing and suggestions.

## Acknowledgements

We want to thank Kristina Haase (EMBL Barcelona) for her support and careful suggestions on the project and manuscript writing, as well as for assistance with the generation of mCherry HBVP with help from Violeta Beltran-Sastre. We acknowledge access to the CRG/UPF flow cytometry facility for mCherry HBVP sorting. We are grateful to Carlota Dobaño (ISGlobal) for access to the Luminex and to Michael Blackman (The Francis Crick Institute) who kindly gifted compound 2. This work was facilitated by the EMBL Electron Microscopy Core Facility (EMCF), with important assistance from Viola Oorschot on sample preparation and processing, and Karel Mocaer and Martin Schorb on image analysis. The great majority of this work was supported by the core program funding of the European Molecular Biology Laboratory (EMBL) and the European Research Council (ERC) under the European Union’s Horizon 2020 research and innovation program (Grant agreement no. 948088). F.K is funded through the Marie Skłodowska-Curie grant agreement (101068552). H.F is supported by a fellowship from the EMBL Interdisciplinary (EI4POD) program under Marie Skłodowska-Curie Actions COFUND (847543). G.M. is supported by RYC 2020–029886 I/ AEI/10.13039/501100011033, co funded by European Social Fund (ESF). ISGlobal received support from the grant CEX2018-000806-S funded by MCIN/AEI/ 10.13039/501100011033, and support from the Generalitat de Catalunya through the CERCA Program. This research is part of the ISGlobal’s Program on the Molecular Mechanisms of Malaria which is partially supported by the Fundación Ramón Areces.

---

**Figure EV1.**
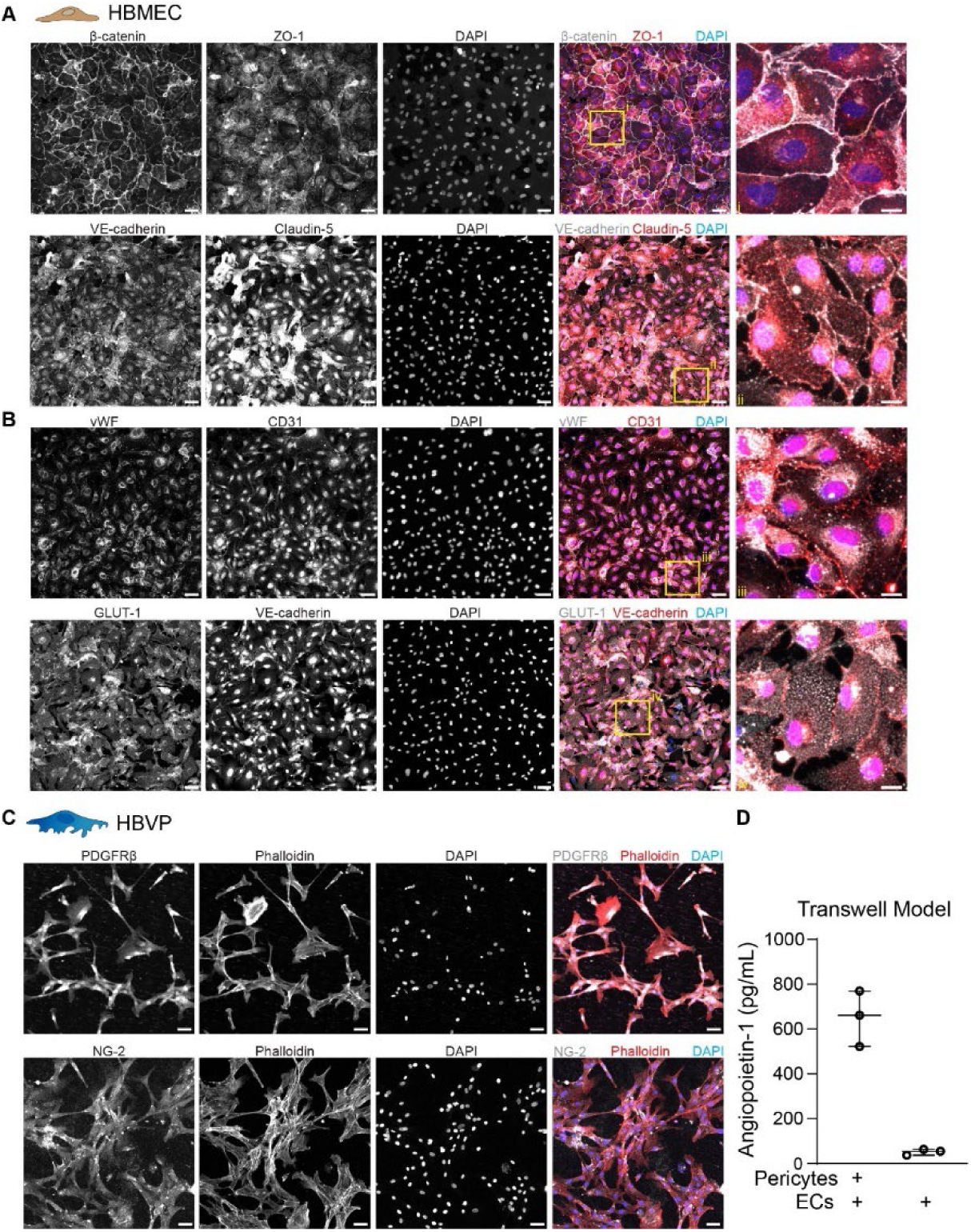
Characterization of brain-specific endothelial and pericyte marker expression and secretion of angiopoietin-Tie axis components. **A** Immunofluorescence maximum z-projection of a 2D HBMEC monolayer stained for adherens and tight junctional markers: β-catenin, VE-cadherin, ZO-1 (top) and Claudin-5 (bottom), and 4’, 6-diamidino-2-phenylindole (DAPI). The merged image includes the adherens junction markers (white), tight junction markers (red) and DAPI labeling (blue). Scale bars: 50 µm and 10 µm for the insets. **B** Immunofluorescence maximum z-projection of a 2D HBMEC monolayer stained for vWF, brain glucose transporter GLUT-1, CD31, VE-cadherin and DAPI. The merge image includes vWF or GLUT-1 (white), CD31 or VE-cadherin (red) and DAPI labeling (blue). Insets in A and B display zoomed merge region of interests (ROIs) to better highlight the cellular localization of each marker. Scale bars: 50 µm and 10 µm for the insets. **C** Immunofluorescence maximum z-projection of a 2D HBVP monolayer stained for PDGFRβ (top) and NG-2 (bottom), Phalloidin and DAPI. The merge staining includes the pericyte markers (white), phalloidin (red) and DAPI labeling (blue). Scale bars: 50 µm. **D** Concentration of secreted angiopoietin-1 in supernatant obtained from either HBVP-HBMEC co-culture or HBMEC-only monolayers grown in a transwell model. (n = 3 independent experiments. Data information: Box and whisker plots display the median and the minimum and maximum data points.

**Figure EV2.**
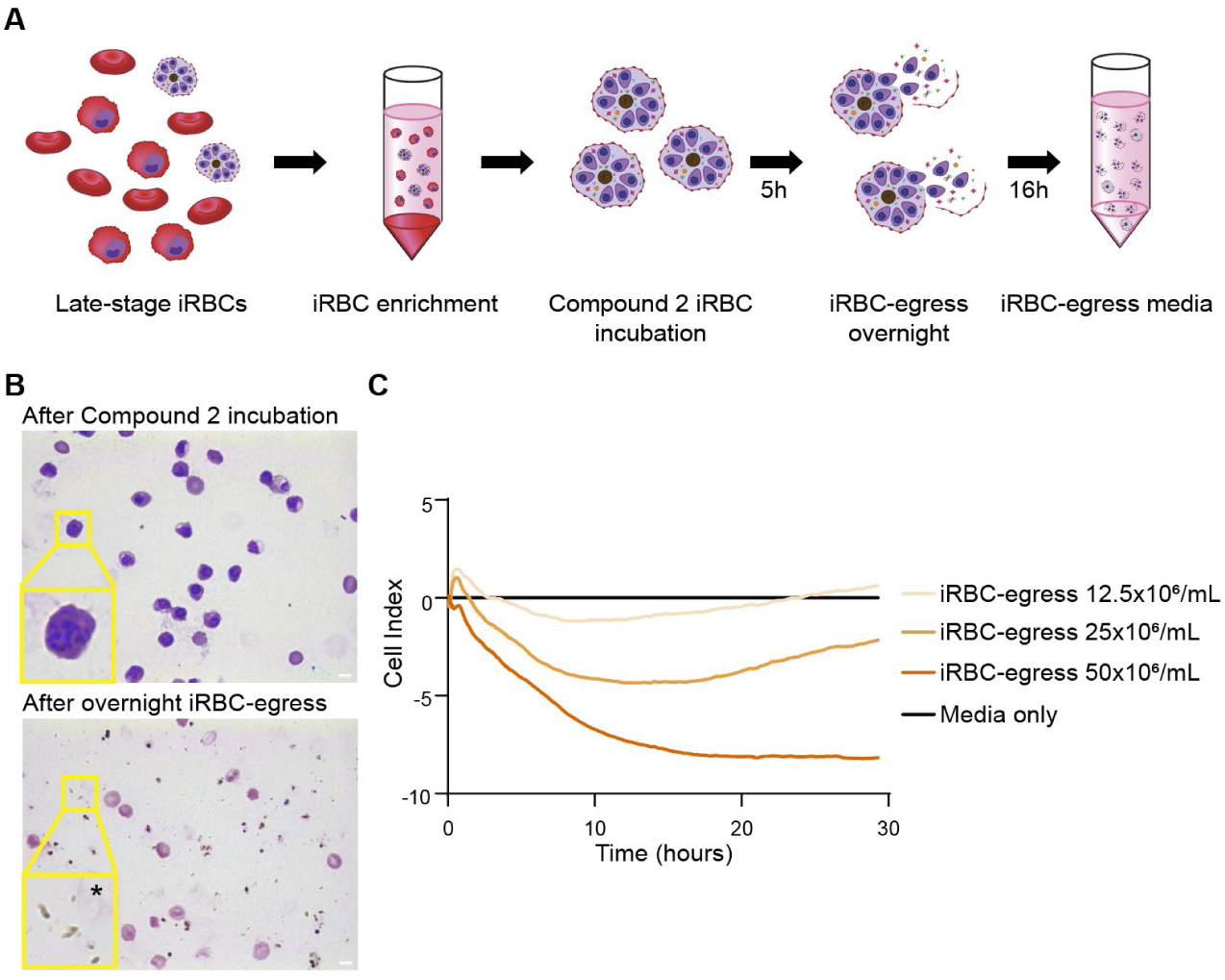
Generation of endothelial barrier disruptive iRBC-egress media. **A** Schematic representation of the protocol to make iRBC-egress media. In short, late-stage *P. falciparum-*iRBC are purified by a gelaspan gradient separation and then incubated for 5 hours with compound-2 to synchronize them at the point of egress. Compound-2 is removed and the *P. falciparum-*iRBC are resuspended in vascular growth media and left overnight on a shaker at 50 rpm to egress. **B** A thin smear of tightly synchronized schizonts before the removal of Compound-2 (yellow inset highlights a ROI with a schizont-stage iRBC) or the resultant iRBC-egress media before centrifugation (yellow inset highlights a ROI with free hemazoin particles next to an iRBC ghost, denoted with an asterisk) stained with giemsa. Scale bars: 5 μm. **C** Representative recording data of xCELLigence measurements on 2D HBMEC monolayers in the absence of HBVP taken after addition of iRBC-egress media at different concentrations. Data is normalized to the media-only control.

**Figure EV3.**
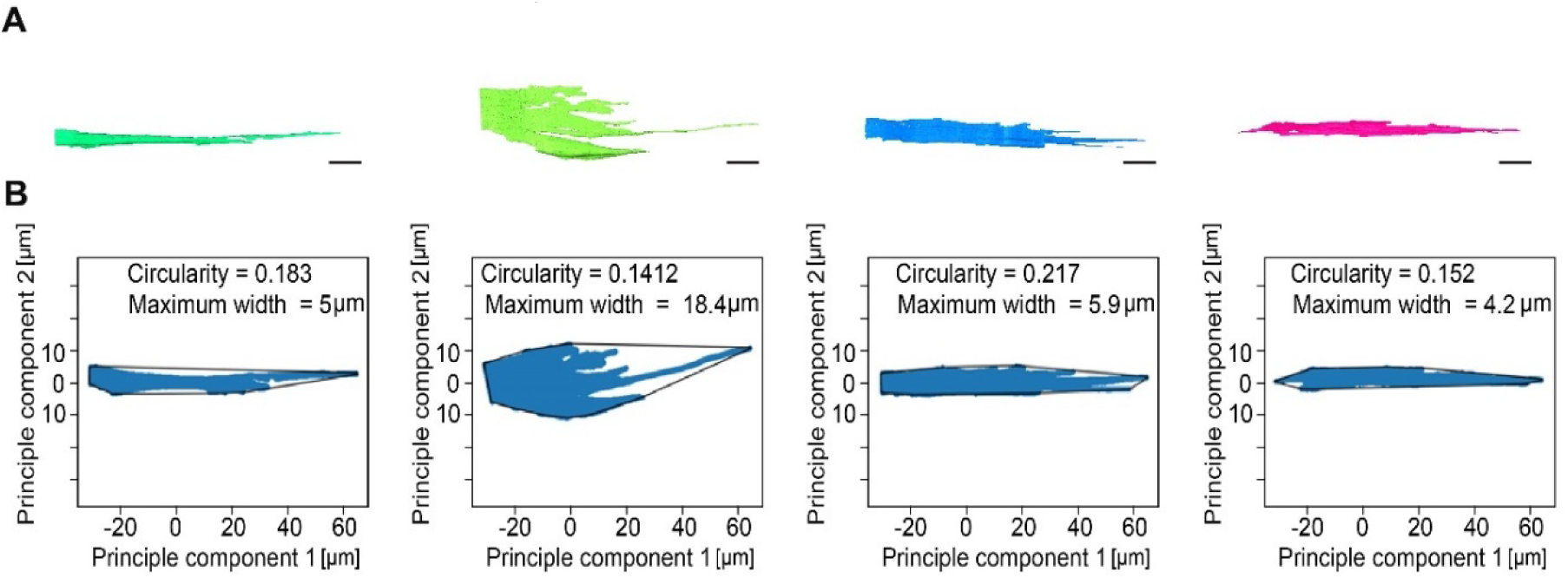
Analysis of pericyte morphological features. **A** The analysis pipeline begins with the extraction of 3D segmented pericyte meshes. Shown here are four representative pericytes. Scale bars: 10 µm. **B** Pericyte geometry is flattened into two principal dimensions using PCA by a Python-based image analysis pipeline. The 2D geometry’s shape is then determined using convex hull analysis. To describe pericyte morphology, pericyte cell borders are analyzed for circularity, calculated as 4π*Area/Perimeter^2^, where a value of 1 indicates a perfect circle and values approaching 0 indicate increasingly elongated shapes, and maximum width.

**Figure EV4.**
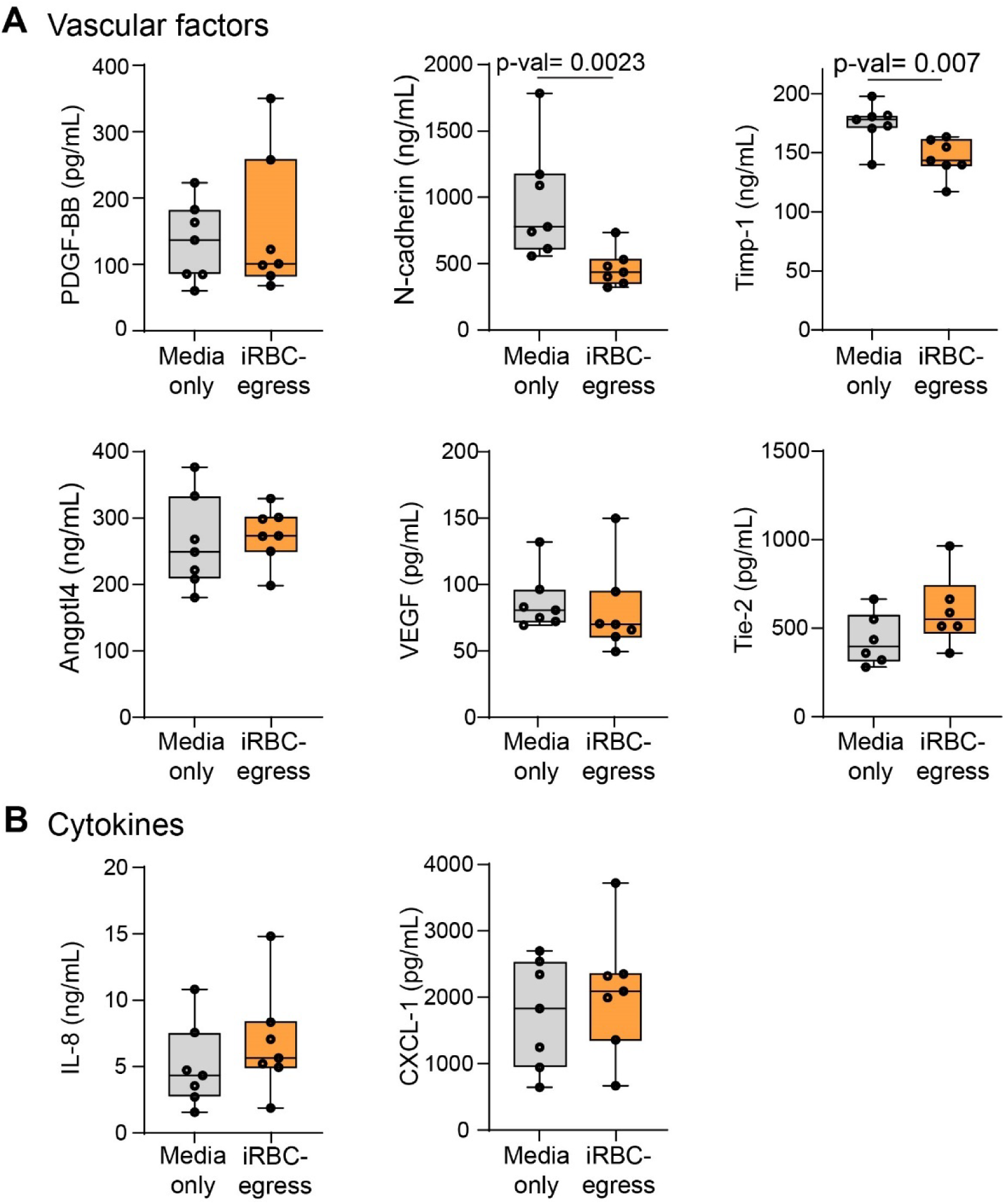
iRBC egress products cause alteration of endothelial cell-pericyte interaction markers. **A** Concentrations of vascular factors PDGF-BB, N-cadherin, Timp-1, Angptl4, VEGF and Tie-2 measured by Luminex from 3D brain microvessels supernatants treated with media only or iRBC-egress media for 18-hours. Box and whisker plots display the median, 25th and 75th percentiles and the minimum and maximum data points. (n_= 7 supernatants pooled from 2-3 devices each). **B** Concentrations of released cytokines IL-8 and CXCL-1 measured by Luminex from 3D brain microvessels supernatants treated with media only or iRBC-egress media for 18-hours. Box and whisker plots display the median, 25th and 75th percentiles and the minimum and maximum data points. (n_= 7 supernatants pooled from 2-3 devices each). Data information: Statistical significance is analyzed by Mann-Whitney U test (**A** and **B**).

**Figure EV5.**
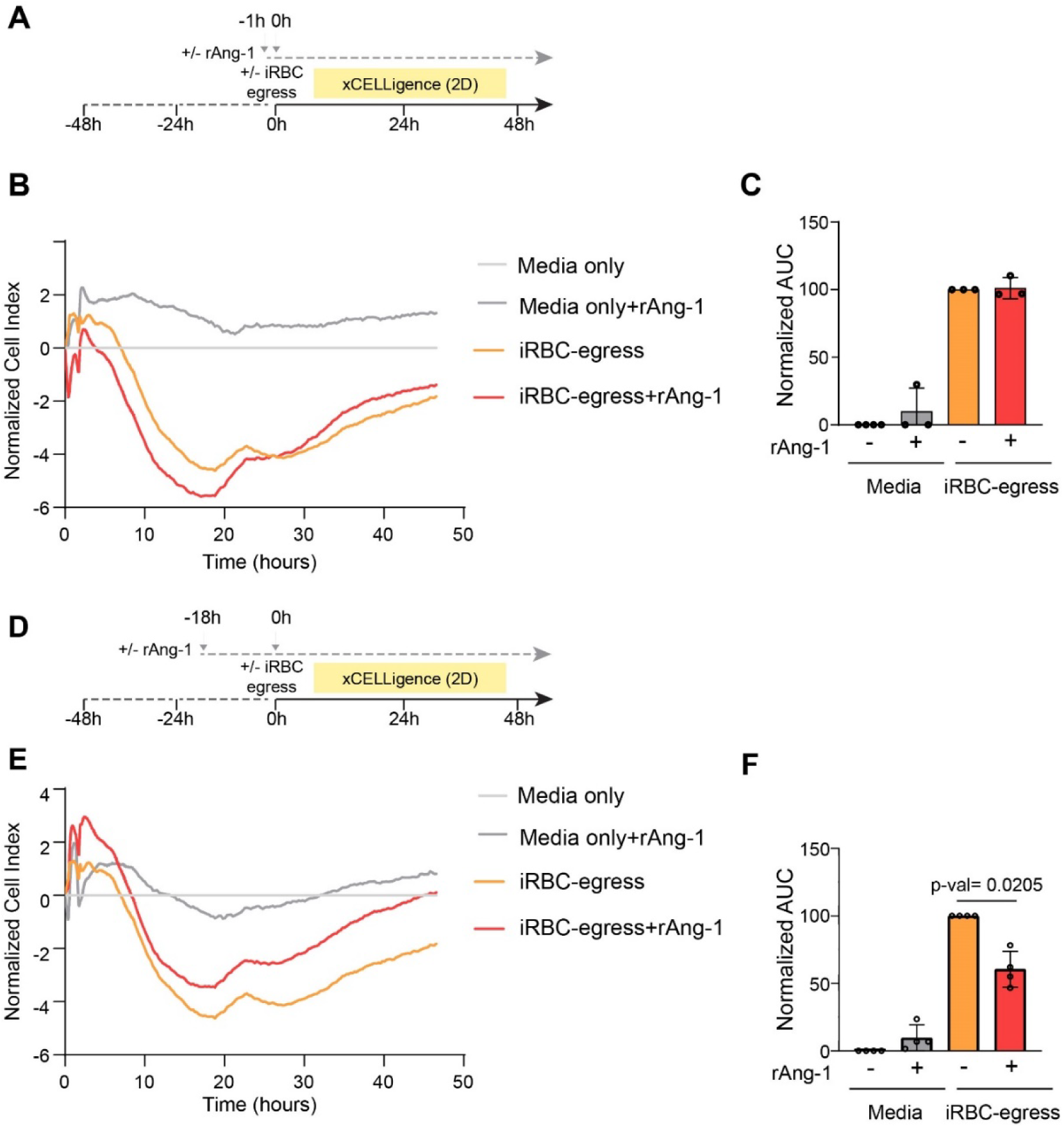
18-hour recombinant Ang-1 pre-treatment partially protects against 2D endothelial monolayer permeability increase induced by iRBC egress products. **A** Experimental outline of short-term (1-hour) incubation with rAng-1 in a 2D HBMEC monolayer in the absence of HBVP. **B** Representative recording data of xCELLigence measurements taken after a 1-hour +/- rAng-1 pre-treatment followed by +/- iRBC-egress media addition. All conditions were normalized to the media only control. **C** Area under the curve analysis of with iRBC-egress media normalized as 100 (n = 3 independent experiments run in triplicate). **D** Experimental outline of long-term (18-hour) incubation with rAng-1 in a 2D HBMEC monolayer in the absence of HBVP. **E** Representative recording data of xCELLigence measurements taken after an 18-hour +/- rAng-1 pre-treatment followed by +/- iRBC-egress media addition. All conditions were normalized to the media only control. **F** Area under the curve analysis of 4 independent experiments run in triplicate with iRBC-egress media normalized as 100. (n = 4 independent experiments run in triplicate). Data information: Error bars represent mean +/- standard deviation. Statistical significance is measured by repeated measures one-way ANOVA test with Dunnett’s multiple comparisons test (**C** and **F**).

**Figure EV6.**
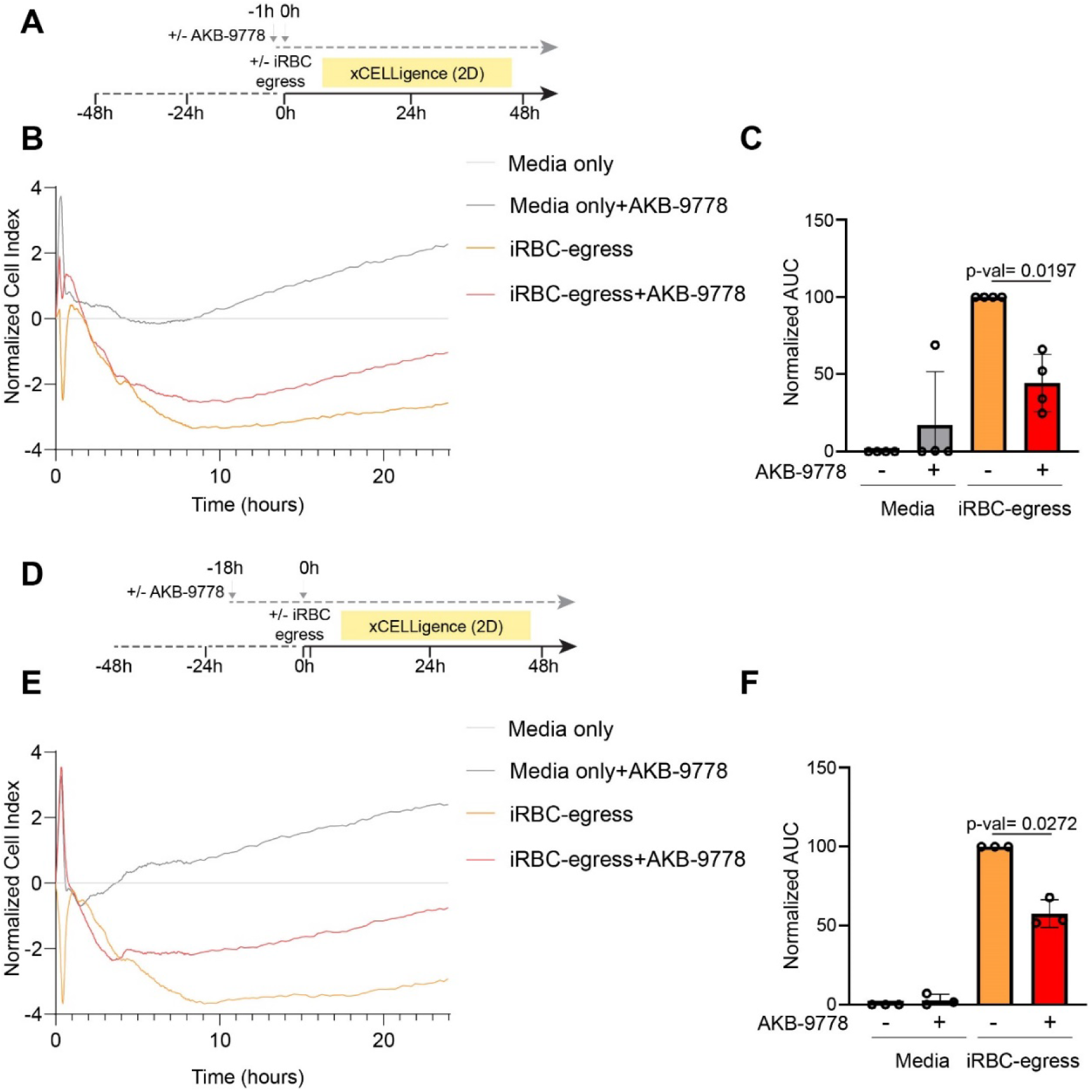
1- and 18-hour AKB-9778 pre-treatment partially protects against 2D endothelial monolayer permeability increase induced by iRBC egress products. **A** Experimental outline of short-term (1-hour) incubation with AKB-9778 in a 2D HBMEC monolayer in the absence of HBVP. **B** Representative recording data of xCELLigence measurements taken after a 1-hour +/- AKB-9778 pre-treatment followed by +/- iRBC-egress media addition. All conditions were normalized to the media only control. **C** Area under the curve analysis with iRBC-egress media normalized as 100 (n= of 4 independent experiments run in triplicate). **D** Experimental outline of long-term (18-hour) incubation with AKB-9778 in a 2D HBMEC monolayer in the absence of HBVP. **E** Representative recording data of xCELLigence measurements taken after an 18-hour +/- rAng-1 pre-treatment followed by +/- iRBC-egress media addition. All conditions were normalized to the media only control. **F** Area under the curve analysis with iRBC-egress media normalized as 100. (n = 3 independent experiments run in triplicate). Data information: Error bars represent mean +/- standard deviation. Statistical significance is measured by repeated measures one-way ANOVA test with Dunnett’s multiple comparisons test (**C** and **F**).

## Notes

https://github.com/waleedmirzaPhD/cell_analysis_toolkit

### Competing Interest Statement

The authors have declared no competing interest.

### Summary of Updates

Figure 5: Addition of AKB-9778 rescue experiment after exposure to P. falciparum egress products

