## Supplemental information for "*Plasmodium falciparum* impairs Ang-1 secretion by pericytes in a 3D brain microvessel model"

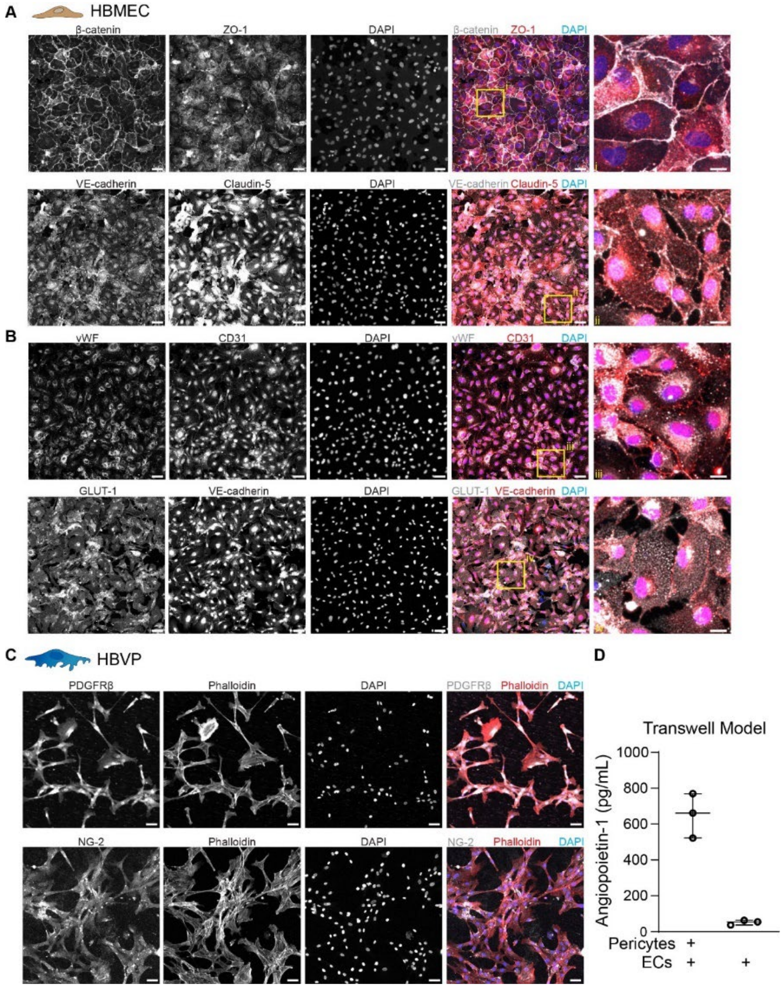


**Figure EV1 - Characterization of brain-specific endothelial and pericyte marker expression and secretion of angiopoietin-Tie axis components.**

**A** Immunofluorescence maximum z-projection of a 2D HBMEC monolayer stained for adherens and tight junctional markers: β-catenin, VE-cadherin, ZO-1 (top) and Claudin-5 (bottom), and 4', 6-diamidino-2-phenylindole (DAPI). The merged image includes the adherens junction markers (white), tight junction markers (red) and DAPI labeling (blue). Scale bars: 50 µm and 10 µm for the insets.

**B** Immunofluorescence maximum z-projection of a 2D HBMEC monolayer stained for vWF, brain glucose transporter GLUT-1, CD31, VE-cadherin and DAPI. The merge image includes vWF or GLUT-1 (white), CD31 or VE-cadherin (red) and DAPI labeling (blue). Insets in A and B display zoomed merge region of interests (ROIs) to better highlight the cellular localization of each marker. Scale bars: 50 µm and 10 µm for the insets.

**C** Immunofluorescence maximum z-projection of a 2D HBVP monolayer stained for PDGFRβ (top) and NG-2 (bottom), Phalloidin and DAPI. The merge staining includes the pericyte markers (white), phalloidin (red) and DAPI labeling (blue). Scale bars: 50 µm.

**D** Concentration of secreted angiopoietin-1 in supernatant obtained from either HBVP-HBMEC co-culture or HBMEC-only monolayers grown in a transwell model. (n = 3 independent experiments.

Data information: Box and whisker plots display the median and the minimum and maximum data points.

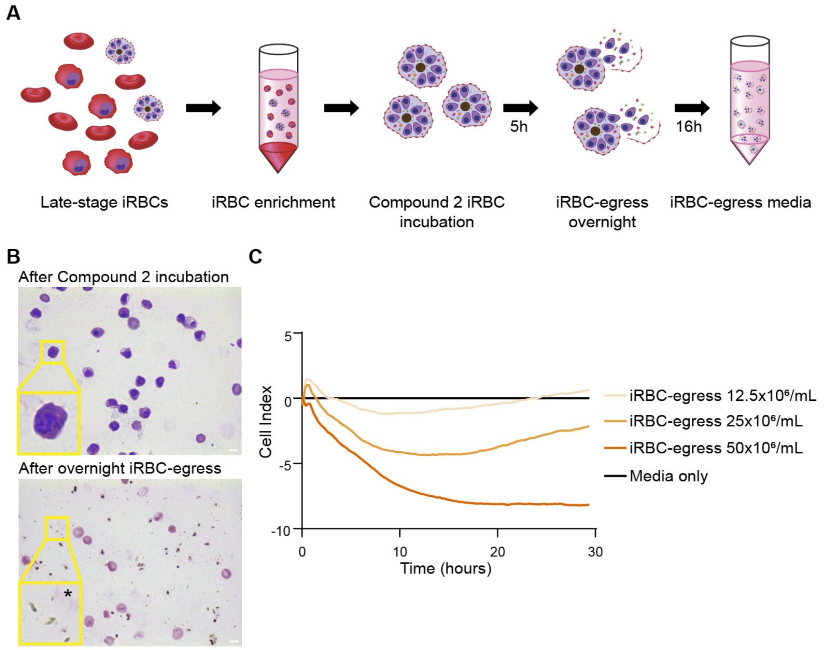


**Figure EV2 - Generation of endothelial barrier disruptive iRBC-egress media.**

**A** Schematic representation of the protocol to make iRBC-egress media. In short, late-stage *P. falciparum*-iRBC are purified by a gelaspan gradient separation and then incubated for 5 hours with compound-2 to synchronize them at the point of egress. Compound-2 is removed and the *P. falciparum*-iRBC are resuspended in vascular growth media and left overnight on a shaker at 50 rpm to egress.

**B** A thin smear of tightly synchronized schizonts before the removal of Compound-2 (yellow inset highlights a ROI with a schizont-stage iRBC) or the resultant iRBC-egress media before centrifugation (yellow inset highlights a ROI with free hemazoin particles next to an iRBC ghost, denoted with an asterisk) stained with giemsa. Scale bars: 5 µm.

**C** Representative recording data of xCELLigence measurements on 2D HBMEC monolayers in the absence of HBVP taken after addition of iRBC-egress media at different concentrations. Data is normalized to the media-only control.

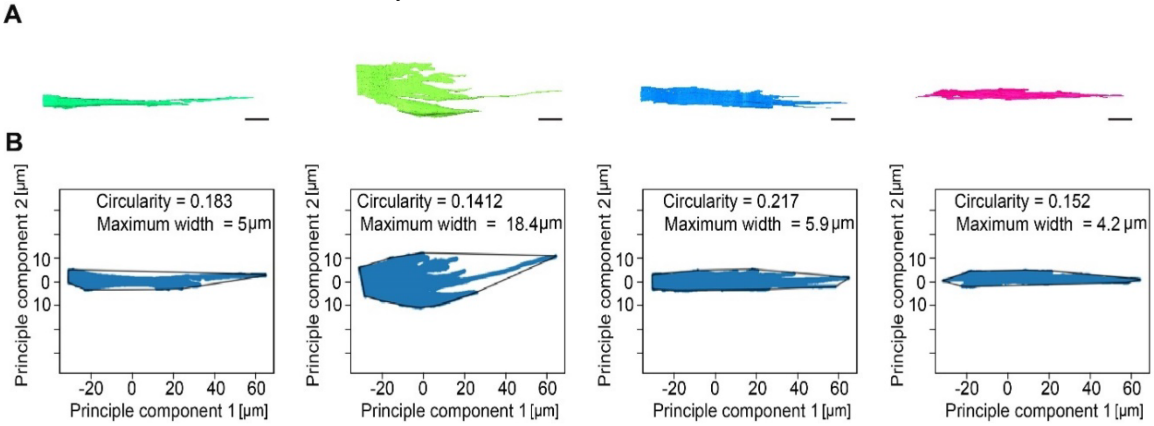


**Figure EV3 - Analysis of pericyte morphological features.**

**A** The analysis pipeline begins with the extraction of 3D segmented pericyte meshes. Shown here are four representative pericytes. Scale bars: 10 µm.

**B** Pericyte geometry is flattened into two principal dimensions using PCA by a Python-based image analysis pipeline. The 2D geometry's shape is then determined using convex hull analysis. To describe pericyte morphology, pericyte cell borders are analyzed for circularity, calculated as $4\pi$*Area/Perimeter$^2$, where a value of 1 indicates a perfect circle and values approaching 0 indicate increasingly elongated shapes, and maximum width.

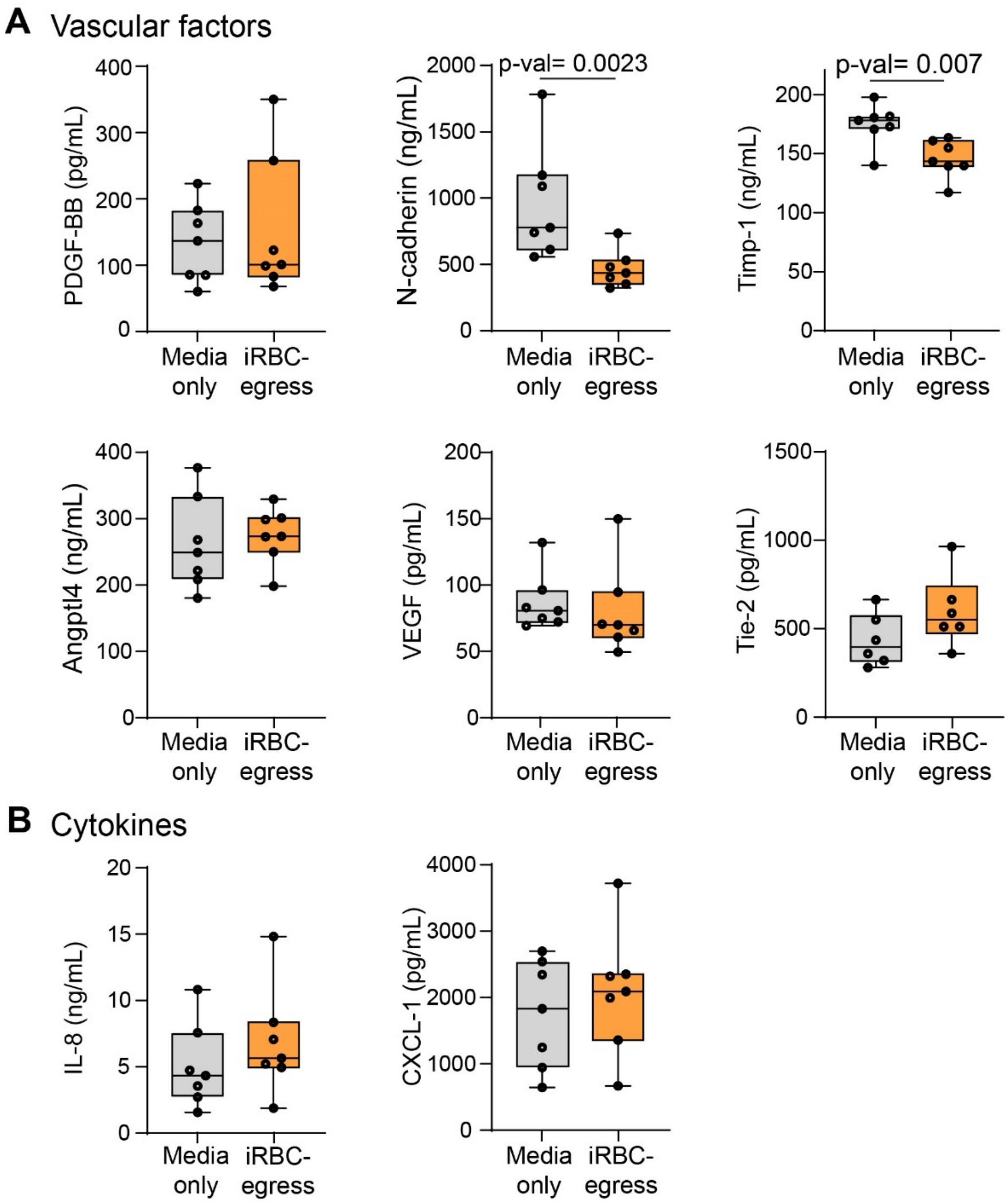


**Figure EV4 - iRBC egress products cause alteration of endothelial cell-pericyte interaction markers**.

**A** Concentrations of vascular factors PDGF-BB, N-cadherin, Timp-1, Angptl4, VEGF and Tie-2 measured by Luminex from 3D brain microvessels supernatants treated with media only or iRBC-egress media for 18-hours. Box and whisker plots display the median, 25th and 75th percentiles and the minimum and maximum data points. (n_= 7 supernatants pooled from 2-3 devices each).

**B** Concentrations of released cytokines IL-8 and CXCL-1 measured by Luminex from 3D brain microvessels supernatants treated with media only or iRBC-egress media for 18-hours. Box and whisker plots display the median, 25th and 75th percentiles and the minimum and maximum data points. (n_= 7 supernatants pooled from 2-3 devices each).

Data information: Statistical significance is analyzed by Mann-Whitney U test (**A** and **B**).

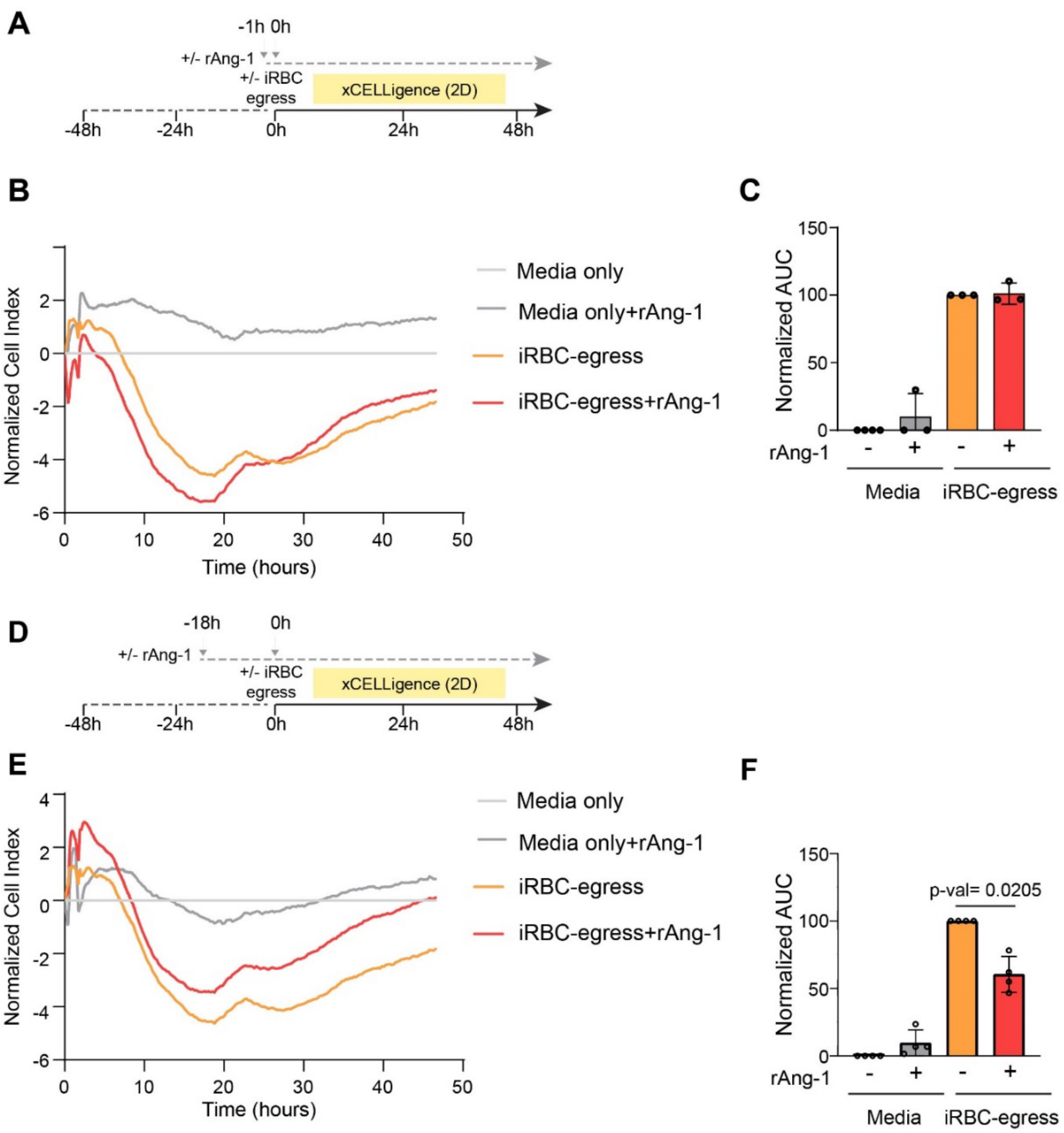


**Figure EV5 - 18-hour recombinant Ang-1 pre-treatment partially protects against 2D endothelial monolayer permeability increase induced by iRBC egress products.**

**A** Experimental outline of short-term (1-hour) incubation with rAng-1 in a 2D HBMEC monolayer in the absence of HBVP.

**B** Representative recording data of xCELLigence measurements taken after a 1-hour +/-rAng-1 pre-treatment followed by +/- iRBC-egress media addition. All conditions were normalized to the media only control.

**C** Area under the curve analysis of with iRBC-egress media normalized as 100 (n = 3 independent experiments run in triplicate).

**D** Experimental outline of long-term (18-hour) incubation with rAng-1 in a 2D HBMEC monolayer in the absence of HBVP.

**E** Representative recording data of xCELLigence measurements taken after an 18-hour +/- rAng-1 pre-treatment followed by +/- iRBC-egress media addition. All conditions were normalized to the media only control.

**F** Area under the curve analysis of 4 independent experiments run in triplicate with iRBC-egress media normalized as 100. (n = 4 independent experiments run in triplicate).

Data information: Error bars represent mean +/- standard deviation. Statistical significance is measured by repeated measures one-way ANOVA test with Dunnett’s multiple comparisons test (**C** and **F**).

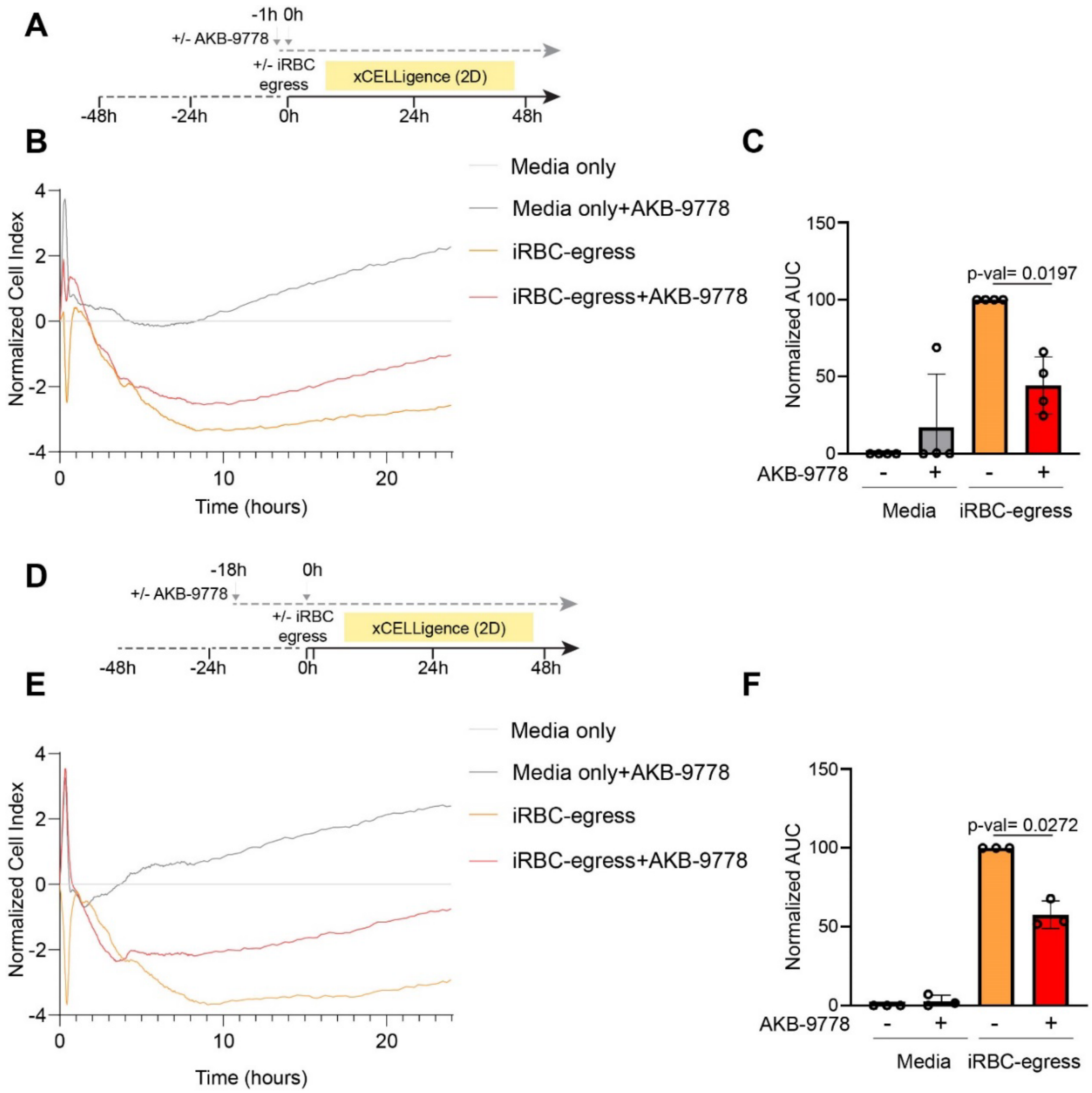


**Figure EV6 - 1- and 18-hour AKB-9778 pre-treatment partially protects against 2D endothelial monolayer permeability increase induced by iRBC egress products.**

**A** Experimental outline of short-term (1-hour) incubation with AKB-9778 in a 2D HBMEC monolayer in the absence of HBVP.

**B** Representative recording data of xCELLigence measurements taken after a 1-hour +/-AKB-9778 pre-treatment followed by +/- iRBC-egress media addition. All conditions were normalized to the media only control.

**C** Area under the curve analysis with iRBC-egress media normalized as 100 (n= of 4 independent experiments run in triplicate).

**D** Experimental outline of long-term (18-hour) incubation with AKB-9778 in a 2D HBMEC monolayer in the absence of HBVP.

**E** Representative recording data of xCELLigence measurements taken after an 18-hour +/- rAng-1 pre-treatment followed by +/- iRBC-egress media addition. All conditions were normalized to the media only control.

**F** Area under the curve analysis with iRBC-egress media normalized as 100. (n = 3 independent experiments run in triplicate).

Data information: Error bars represent mean +/- standard deviation. Statistical significance is measured by repeated measures one-way ANOVA test with Dunnett's multiple comparisons test (**C** and **F**).
